# Polarized anionic phospholipids and exocytosis are implicated in the polarized recruitment of budding yeast AP180, an endocytic initiator

**DOI:** 10.1101/2024.10.08.617284

**Authors:** Paul Marchando, Gean Hu, Feng Yuan, Jordan M. Ngo, Yidi Sun, David G. Drubin

## Abstract

Understanding of the mechanisms that initiate clathrin-mediated endocytosis (CME) is incomplete. Recent studies in budding yeast identified the endocytic adaptor proteins Yap1801/Yap1802 (budding yeast AP180) as key CME factors that promote CME initiation in daughter cells during polarized growth, but how Yap1801/2 are recruited preferentially to the plasma membrane of daughter cells is not clear. The only known cargos for Yap1801/2 in yeast are the synaptobrevins Snc1 and Snc2, which serve as v-SNARES for exocytic vesicles reaching the plasma membrane and are crucial for polarized cell growth. In this study, we examine the spatiotemporal dynamics of functional, fluorescent protein-tagged Snc2 expressed from its endogenous locus and provide evidence that, along with anionic phospholipids, Snc2 specifically recruits Yap1802 to growing daughter cells. This protein-protein interaction creates a direct link between polarized secretion and CME and has further implications in CME initiation.

## Introduction

While much research has focused on how clathrin-mediated endocytosis (CME) is initiated, consensus about the molecular mechanism has not been reached (Godlee and Kaksonen, 2013). Rab GTPases are known to initiate many trafficking processes, but no single trigger or pioneering factor for CME has been identified (Kreis *et al*., 1995; Godlee and Kaksonen, 2013; Martin and Arkowitz, 2014), although membrane-bound cargo has been proposed to play a role (Taylor *et al*., 2011; Goode *et al*., 2015; Pedersen *et al*., 2020).

In the budding yeast *Saccharomyces cerevisiae*, recent studies demonstrated that CME site initiation and maturation dynamics depend on position on the plasma membrane within a growing, polarized yeast cell (Layton *et al*., 2011; Pedersen *et al*., 2020; Sun *et al*., 2024). Budding yeast proliferate by forming daughter cells, also referred to as “buds,” on the surface of a mother cell. Exocytosis is highly biased toward the daughter cell, so molecules delivered to the plasma membrane during exocytosis, such as SNARE proteins, are enriched near these exocytic sites (Field and Schekman, 1980). Since CME is the major pathway that retrieves material from the plasma membrane back into the cell interior, the presence of cargo concentrated in the bud might affect CME site initiation or maturation (Carroll *et al*., 2012; Cocucci *et al*., 2012; Brach *et al*., 2014). Indirect approaches have supported this possibility, but a direct link between polarized secretion and polarized CME (originally observed as polarized actin patches) has yet to be demonstrated (Adams and Pringle, 1984; Goode *et al*., 2015; Pedersen *et al*., 2020).

CME in budding yeast displays two temporal phases with distinct behaviors (Pedersen *et al*., 2020). The transition between these phases coincides with differential Pan1 (budding yeast intersectin homolog) behavior in mother and daughter cells (Sun *et al*., 2024). Pan1 undergoes a “low-abundance” phase followed by a “high-abundance” phase in mother cells, while only displaying the high-abundance phase in daughters (Sun *et al*., 2024). We thus define early-arriving proteins as those that appear at CME sites before the high-abundance Pan1 phase, and late-arriving proteins as those that appear during or after the high-abundance Pan1 phase (Brach *et al*., 2014; Bradford *et al*., 2015; Sun *et al*., 2015, 2017, 2024; Pedersen *et al*., 2020). Early-arriving proteins show position-dependent differences in lifetimes (short and regular in daughters, longer and variable in mothers), while late-arriving proteins exhibit more regular lifetimes regardless of cell growth stage or position within the cell (Pedersen *et al*., 2020; Sun *et al*., 2024).

This heterogeneity in CME site behavior is partly driven by the endocytic adaptor complex AP180 (Yap1801 and Yap1802 in yeast, abbreviated as Yap1801/2) (Sun *et al*., 2024). Yap1801/2 interact cooperatively with the FCHo1/2 homolog Syp1 to recruit the EPS15 homolog Ede1 to the bud cortex, initiating the early stages of CME (Lu and Drubin, 2017; Sun *et al*., 2024). Yap1801/2 are polarized strongly to the bud during yeast cell growth while mother cells have nearly undetectable Yap1801/2, explaining the observed position-dependent differences in CME initiation rates and lifetimes (Sun *et al*., 2024). While Yap1801/2’s role in CME initiation as perhaps the earliest acting factor is more clear, the factors upstream of Yap1801/2 recruitment remain speculative (Sun *et al*., 2024).

Yap1801/2 have one known cargo, the v-SNAREs Snc1 and Snc2 (abbreviated as Snc1/2), homologs of the mammalian synaptobrevins, which depend of CME for their retrieval from the plasma membrane (Burston *et al*., 2009; Miller *et al*., 2011). Snc1/2 are the only known v-SNAREs in budding yeast that facilitate the fusion of secretory vesicles with the plasma membrane (Burri and Lithgow, 2004). In addition to binding Snc1/2, Yap1801/2 can directly bind anionic lipid species (Ford *et al*., 2001). Since both Snc1/2 and anionic lipid species, particularly phosphatidylserine, are enriched in daughter cells, Snc1/2 and anionic lipids are poised to play a role in the asymmetrical behavior of Yap1801/2 in yeast undergoing bud growth (Burston *et al*., 2009; Sun and Drubin, 2012; Meca *et al*., 2019). The residues in Yap1801/2 mediating both binding activities are well characterized, and precise perturbation of both Snc1/2 and lipid binding could reveal if both are equally important for Yap1802 recruitment or if one overshadows the other.

The yeast synaptobrevins Snc1/2 thus provide a unique opportunity to study the earliest stages of CME and better characterize a yeast CME cargo, a task that has remained difficult and yielded mixed results in past studies (Toshima *et al*., 2006; Carroll *et al*., 2009). Using live-cell microscopy, we compare the behaviors of fluorescent Snc1 and Snc2 expressed from their natural chromosomal loci and provide the first spatiotemporal analysis of Snc2 expressed at endogenous levels. This analysis highlights a complex relationship between Snc2 and CME, which varies depending on the cell growth stage. Additional experiments show that Snc2 and anionic phospholipids collaborate to recruit the endocytic adaptor Yap1802 to daughter cells during growth, linking polarized secretion to CME initiation.

## Results and Discussion

### Functional GFP-tagged Snc2 fusion protein expressed from its endogenous genomic locus displays complex spatiotemporal behaviors relative to the CME machinery

To analyze the spatial distribution of Snc1 and Snc2 in yeast cells, we created in-frame fusions of the corresponding genes at their endogenous genomic loci with sequences encoding GFP and visualized the N-terminally-tagged fluorescent fusion proteins in cells by Airyscan (Zeiss) confocal microscopy (Figure 1A, Figure S1A, arrows). The two proteins displayed similar subcellular localization, appearing preferentially in growing daughter cells, in agreement with previous studies of overexpressed Snc1, and they both decorated internal structures that are presumably membrane-bound organelles (Gurunathan *et al*., 2000, 2002; Lewis *et al*., 2000; Valdez-Taubas and Pelham, 2003; Burston *et al*., 2009). The GFP-Snc1 fusions displayed weak signals, consistent with the low signal observed on immunoblots, suggesting a greater abundance of Snc2 (Figure S2A). In addition, *snc2Δ* strains were more severely impaired in growth than *snc1Δ* strains (Figure S2B). *SNC2* transcripts outnumber *SNC1* transcripts by nearly threefold, perhaps explaining the differential behaviors even though the paralogs display 76% sequence identity (Katz *et al*., 1998; Grote *et al*., 2000; Gurunathan *et al*., 2000; Lipson *et al*., 2009). Due to the greater abundance and importance of Snc2 for cell growth, we focus on GFP-Snc2 henceforth. Since GFP-Snc2 expressed from the endogenous promoter was able to rescue the growth defects of *snc2Δ* cells, we conclude that this fusion protein is functional and likely to serve as a faithful reporter of the spatiotemporal dynamics of these v-SNARES (Figure S2B).

**FIGURE 1:**
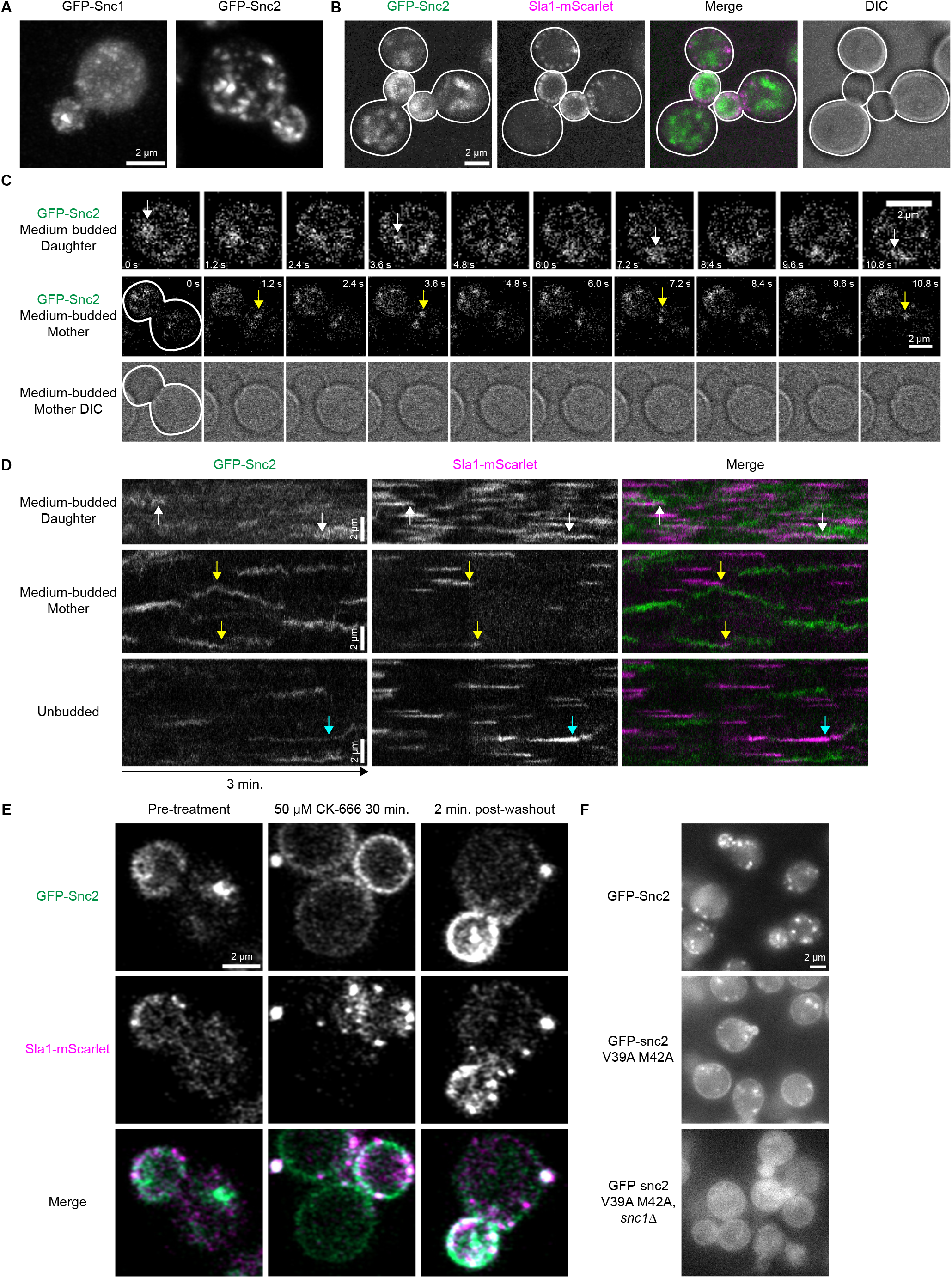
Spatiotemporal localization of fluorescent protein-tagged Snc1 and Snc2 expressed from their endogenous genomic loci in normal, drug-treated, and mutant cells. **A**. Maximum intensity projections of Airyscan confocal z-stacks of fixed yeast cells expressing GFP-Snc1 (with intron removed from the *SNC1* gene) and GFP-Snc2, respectively. **B**. Maximum intensity projections from a time-lapse movie of cells expressing GFP-Snc2 and Sla1-mScarlet acquired using a Nikon confocal microscope equipped with an NSPARC detector (see Materials and Methods). Here two budded cells and one unbudded cell are outlined. **C**. Montages of time-lapse GFP-Snc2 live-cell NSPARC movies at different stages of cell growth. First row is an expanded image of the bud in the second row. Time interval between each frame is 1.2 seconds. White and yellow arrows in the first and second rows identify unique translating Snc2-positive puncta in daughter and mother cells, respectively. **D**. Circumferential kymographs made from 3-minute NSPARC movies of cells expressing GFP-Snc2 and Sla1-mScarlet in different cell regions and stages of cell growth. White, yellow, and cyan arrows denote unique Snc2/Sla1 interaction events in daughter, mother, and unbudded cells, respectively. Temporal resolution is ∼0.6 seconds. **E**. Still frames from time-lapse Airyscan confocal microscopy movies of wild-type cells expressing GFP-Snc2 and treated with 50 µM CK-666 for 30 minutes, or 2 minutes after washing out the inhibitor (cells in each column are different). **F**. Still frames from widefield time-lapse movies of cells carrying various mutant GFP-Snc2 alleles.

We next studied the complex behaviors and interactions of Snc2 with the CME machinery (reported by Sla1-mScarlet) in live cells using Nikon Spatial Array Confocal (NSPARC) microscopy at sub-second temporal resolution (Supplemental Videos 1 and 2) (Delattre, 2023). Snc2 and Sla1 both showed preferential enrichment at the plasma membrane in daughter cells (Figure 1B, Figure S1B, arrows). Snc2 displayed a mixture of diffuse and punctate signals while Sla1 appeared solely in puncta (Figure 1A, B, Figure S1A, B). Bright Snc2-positive compartments, possibly early endosomes, translated across large lengths of budded daughter cells in several seconds, moving from the plasma membrane through the cell interior and returning to the plasma membrane surface (the same cell and event is depicted in Figure 1C, white arrows, and Supplemental Video 1, arrows). This type of motion is consistent with the known Snc2 retrograde transport to the trans-Golgi network (TGN), fusion with endosomes, and delivery back to the plasma membrane (Gurunathan *et al*., 2000; Lewis *et al*., 2000; Burri and Lithgow, 2004; Best *et al*., 2020). Additionally, punctate cytoplasmic structures moved directionally from mother cells toward daughter cells, consistent with their participation in polarized secretion (Figure 1C, yellow arrows, Supplemental Video 1, arrows).

Using circumferential kymographs created by tracing the outline of the plasma membrane on the vertical axis and projecting the outline through time on the horizontal axis, we better observed how Snc2-positive compartments interact with the plasma membrane and the CME machinery in mother vs. daughter cells (Figure 1D, Figure S1C-E) (Lu and Drubin, 2017; Sun *et al*., 2017). In daughter cells, GFP-Snc2 appeared on the plasma membrane diffusely, highlighted by the background haziness observed throughout kymographs in Figure 1D (top row) and Figure S1C when compared to other growth stages. Within these kymographs, bright traces of Snc2-positive compartments appeared and disappeared (Figure 1D, white arrows, Figure S1C), coinciding with the Snc2-positive compartments observed in Figure 1C and Supplemental Video 1 that transiently interacted with the plasma membrane. While these bright Snc2-positive traces did generally overlap with Sla1 traces (Figure 1D, white arrows, Figure S1C), the high Snc2 background precludes any direct functional interpretations. In kymographs of mother cells, Snc2 puncta translated laterally across the plasma membrane, occasionally coinciding with the internalization of endocytic vesicles (Figure 1D, yellow arrows, Figure S1D). Unbudded cells displayed a mixture of these behaviors, with brighter traces that fluctuate less in intensity and show longer lifetimes (Figure 1D, third row, Figure S1E). On rare occasions, Snc2-positive punctate traces were observed to track and disappear with Sla1 in a manner consistent with CME cargo behavior (Figure 1D, cyan arrows).

CME perturbation highlights a clear dependency of Snc2 on CME for internalization. Treatment of yeast cells with the Arp2/3 complex inhibitor CK-666 halted endocytic internalization (Hetrick *et al*., 2013) and trapped Snc2 preferentially on the plasma membrane in daughter cells while Snc2 cytoplasmic puncta disappeared (Figure 1E). In these cells, stalled endocytic sites, marked by Sla1-mScarlet, appeared as stationary puncta on the plasma membrane, although Snc2 signal was not noticeably enriched at these puncta. Upon CK-666 washout, Snc2 could again be seen on cytoplasmic puncta and appeared brighter, perhaps due to the drug-induced local Snc2 enrichment in daughter cells (Figure 1E). Using NSPARC microscopy to observe the restart of CME following CK-666 washout with greater spatial and temporal resolution, we more clearly visualized the direct interaction of Snc2 and Sla1-positive traces (Figure S1F). Kymographs of videos acquired directly following CK-666 washout showed increased Snc2/Sla1 colocalization and coincident internalization when compared to unperturbed cells (Figure S1F, white arrows). These kymographs additionally suggested a slight enrichment of Snc2 at stalled Sla1 puncta, demonstrated by the colocalizing parallel traces observed before CME resumes (Figure S1F, yellow arrows). These observations suggest that, in daughter cells, Snc2 tracks and internalizes with the CME machinery like the other canonical CME components, but the superimposed background signal from other stages of Snc2’s complex recycling cycle obscures this behavior at steady state. Observing cargo associated with CME sites is notoriously difficult due to signal-to-noise challenges as cargo saturate CME sites and accumulate on the plasma membrane.

These combined results suggest a complex interplay between the dynamic Snc2 life cycle and CME. Our time-lapse data are consistent with the known functions of Snc2 in polarized secretion and in retrograde transport of endosomes back to the TGN, but the high-speed dynamics of these behaviors and the coincident endocytic internalization of Snc2 with Sla1 are newly reported here. Of particular interest is the observed interaction of cytosolic Snc2-positive compartments with plasma membrane-bound Sla1 traces. These Snc2-positive puncta appear to move to and from the plasma membrane in a coordinated manner, as if collecting newly budded endosomes from CME sites (Figure 1D, yellow arrows, Supplemental Video 2, arrows). This “vacuum cleaner” behavior has also been observed between labeled endosomes and endocytic sites in our previous studies using both fluorescent alpha factor endocytic cargo (Toshima *et al*., 2006) and the lipophilic dye FM 4-64 (Cortesio *et al*., 2015). These events could represent endosomes that fuse with nascent endocytic vesicles via Snc2’s SNARE activity, but whether SNARE function plays an active role in this behavior remains to be investigated. Nonetheless, this behavior suggests a high level of spatial and temporal coordination between CME vesicle internalization and subsequent fusion with early endosomes, and we believe the strains created here can aid further investigation of this mechanism.

### Snc2 internalization from the cell surface via CME is important for cell viability

Previous work on the mammalian synaptobrevins identified residues that mediate their interaction with the endocytic adapter CALM, a mammalian homolog of Yap1801/2 (Grote *et al*., 2000; Gurunathan *et al*., 2000; Miller *et al*., 2011). The residues important for this interaction correspond to V39 and M42 on Snc2. Alanine substitutions at these residues did not affect Snc2 expression in both *SNC1* and *snc1Δ* backgrounds (Figure S2C) but resulted in diffuse Snc2 signal on the plasma membrane in both daughter and mother cells, consistent with an internalization defect and similar to the phenotype observed when endocytosis is stalled with CK-666 (Figure 1F) (Grote *et al*., 2000; Shen *et al*., 2013). This result suggests that the V39A M42A Snc2 mutant is still competent to mediate exocytosis, Snc1/2’s essential function, and that CME of more lowly-expressed Snc1 alone is sufficient for yeast to remain healthy in standard conditions. We tested the latter conclusion in an *snc1Δ* background, wherein the Snc2 mutant again showed diffuse plasma membrane staining in mother and daughter cells, but these cells displayed a severe growth defect, irregular morphologies, and diffuse Snc2 cytoplasmic signal consistent with vacuolar fragmenting or sorting defects observed in unhealthy cells (Figure 1F, S2B) (Grote *et al*., 2000; Stauffer and Powers, 2015). These observations differ from previous findings, which show reduced Snc2 M42A protein levels by immunoblot compared to wild-type Snc2, and no growth defect in snc1Δ snc2 M42A cells. This result suggests that the V39A M42A Snc2 mutant expressed natively shows decreased Snc2 protein levels but is not harmful in an snc1Δ background (Grote *et al*., 2000). This discrepancy could be explained by the prior introduction of the *SNC2* coding sequence, with 500 flanking base pairs on either side of the open reading frame, at the *LEU2* locus, rather than at the native chromosomal locus (Grote *et al*., 2000). Perhaps there are regulatory sequences beyond these 500 base pairs at the native locus that impact *SNC2* expression. Given our observed roughly 7-fold difference in Snc1 and Snc2 protein abundance but only 3-fold difference in RNA expression, there may be differences in the expression and translation of these native loci and transcripts that have yet to be uncovered (Figure S2A) (Lipson *et al*., 2009).

These observations indicate that CME of some Snc1/2 molecules is essential for normal cell physiology. While it is possible that the V39A M42A mutation impacts other cellular processes in which Snc2 is involved, the conserved methionine residue has been shown to specifically mediate the interaction of synaptobrevins with CALM/AP180, and previous studies demonstrated that CME of SNAREs and their ability to mediate exocytosis are separable activities (Miller *et al*., 2011; Ma and Burd, 2019).

### Roles for polarized anionic phospholipid and Snc2 localization in polarized Yap1802 recruitment

We next created specific point mutations in Yap1802 to investigate the relationship between Yap1802, Snc2, and negatively charged lipid species during polarized growth. Residues of the mammalian Yap1802 homolog CALM that mediate its interaction with anionic phospholipids and synaptobrevins have been characterized and fall within its ANTH (AP180 N-terminal homology) domain (Ford *et al*., 2001; Miller *et al*., 2011; Pashkova *et al*., 2021). The analogous residues in *S. cerevisiae* Yap1802 correspond to lysines 21 and 23 for lipid interaction and leucine 203 for Snc2 interaction (Figure 2A, B) (Ford *et al*., 2001; Sun *et al*., 2005; Miller *et al*., 2011; Pashkova *et al*., 2021). Alphafold2 multimer analysis strengthened confidence that L203 in Yap1802 interacts with M42 in Snc2 (Figure 2B) (Mirdita *et al*., 2022). Using a CRISPR/Cas9-based approach to generate yap1802 K21E K23E and L203S mutants, we tested whether interactions of Yap1802 with anionic phospholipids and plasma membrane-bound Snc2 might facilitate Yap1802 recruitment to growing daughter cells (Anand *et al*., 2017). Given the high degree of homology between Yap1801 and Yap1802, the observation that Yap1802 is functionally sufficient in *yap1801Δ* cells, and the strong polarization of Yap1802 to small- and medium-sized daughter cells, we decided to conduct experiments on point-mutated Yap1802 mutants in a *yap1801Δ* background (Sun *et al*., 2024). The mutated strains showed no detectable difference in Yap1802 expression levels and did not cause an observable temperature-sensitive cell viability defect (Figure S2D, E). Liposome flotation assays confirmed a lipid binding defect for the yap1802 K21E K23E ANTH domain, which is evident at various concentrations of the mutant YAP1802 ANTH domain compared to the wild-type YAP1802 ANTH domain (Figure S2F).

**FIGURE 2:**
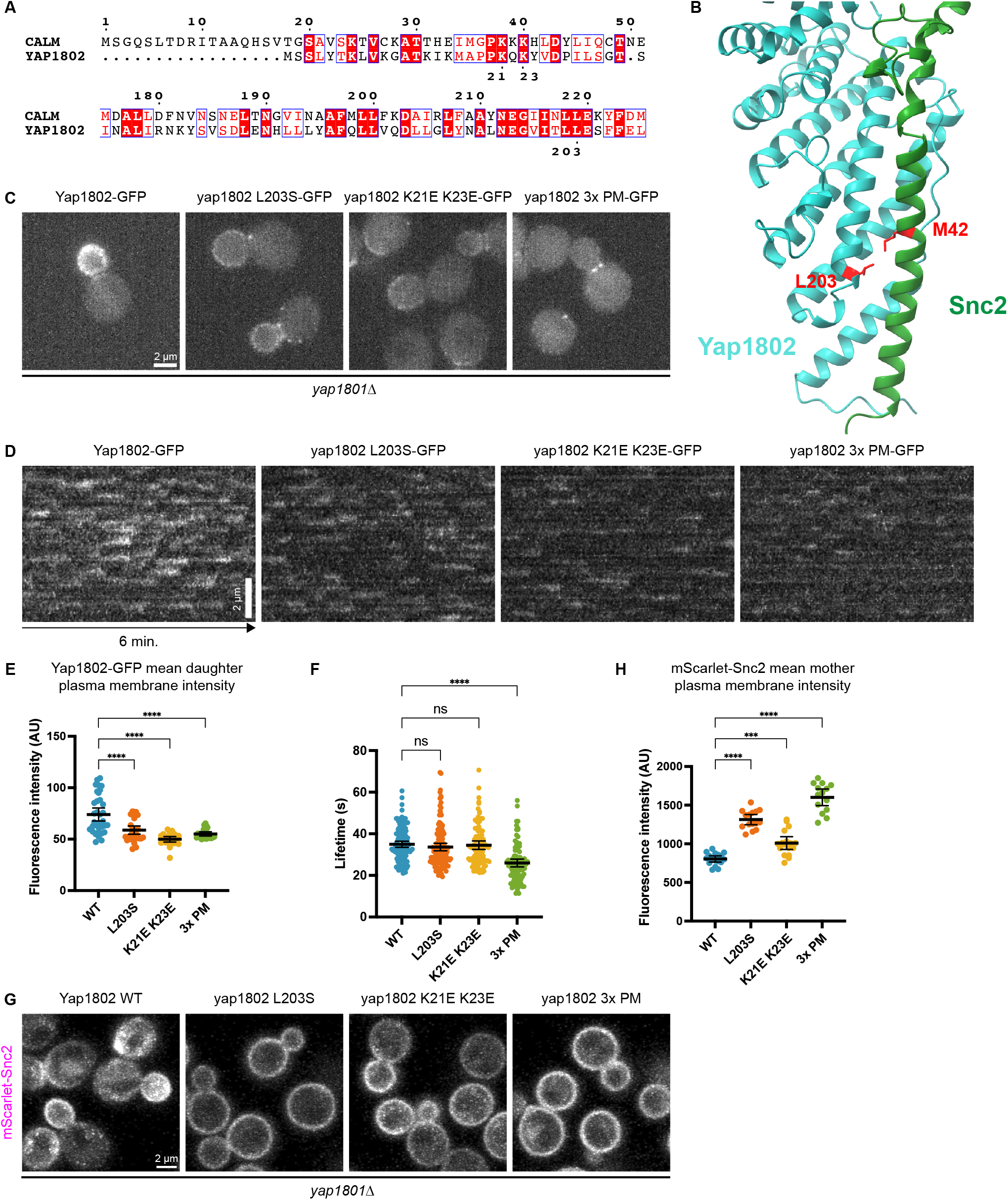
Yap1801/2 is recruited to the daughter cortex by Snc1/2 and negatively charged lipids. **A**. Sequence alignment for human CALM and *S. cerevisiae* Yap1802. Red fill indicates direct matches, and red characters indicate shared residue characteristics (Robert and Gouet, 2014). **B**. Alphafold2 multimer analysis of the ANTH domain of Yap1802 and Snc2, with key residues highlighted in red. **C**. Maximum intensity projections from 6-minute widefield microscopy movies of cells expressing various Yap1802 wild-type or point-mutant alleles, 3x PM refers to the K21E K23E L203S triple point mutant. **D**. Circumferential kymographs made from 6-minute widefield microscopy movies of Yap1802-GFP point mutants in medium-budded daughter cells. Temporal resolution is 2 seconds. **E**. Mean intensities of Yap1802-GFP point mutant line profiles from daughter cell cortexes as in **C**. Right to left, n = 35, 32, 23, 22 **F**. Quantification of Yap1802-GFP point mutant lifetimes. **G**. Maximum intensity projections from 3-minute Airyscan confocal microscopy time-lapse movies of cells expressing wild-type Yap1802 or point mutants and mScarlet-Snc2. Temporal resolution of movies was ∼3.7 seconds. Right to left, n = 127, 129, 106, 97. **H**. Mean intensities of mScarlet-Snc2 line profiles from mother cell cortexes as in **G**. Right to left, n = 17, 15, 17, 14. Groups in **E, F**, and **H** were compared using one-way ANOVA. Center and error bars are mean and 95% confidence interval, *** indicates 0.001 > p > 0.0001, and **** indicates p < 0.0001.

Yeast strains individually expressing the yap1802 K21E K23E mutant (predicted to be defective in lipid binding), the yap1802 L203S mutant (predicted to be defective in Snc2 binding), or the yap1802 3x PM (K21E K23E L203S, predicted to be defective in both lipid and Snc2 binding) each displayed a Yap1802 recruitment defect, which can be best visualized using maximum intensity projections of time-lapse movies (Figure 2C). In these projections, quantification of the mean cortex fluorescence intensity in daughter cells showed a decrease in plasma membrane Yap1802 signal among all point mutants tested (Figure 2E). Additionally, looking closer at these cortex intensity traces in daughter cells revealed a unimodal curve with a maximum near the middle of the trace, coinciding with the very tip of the growing bud (Figure S3A). Each individual mutant lost this curved Yap1802 trace and showed reduced intensity along the length of each trace. Individual Yap1802 endocytic events in budded cells were still observed in each mutant, with the lipid and Snc2 mutants maintaining normal lifetimes compared to cells expressing wild-type Yap1802 (Figure 2D, 2F). However, the lifetimes of yap1802 3x PM-GFP events were significantly shorter than those for the wild-type cells and individual mutants (Figure 2F). These 3x PM lifetimes are consistent with those observed in strains that lack the ability to localize Ede1 properly, such as *yap1801Δ/yap1802Δ* double null mutants and *ede1Δ* cells (Brach *et al*., 2014; Sun *et al*., 2024). Since Yap1802 is required for proper Ede1 localization, this observation suggests that a major part of Yap1802’s membrane-binding ability is contained within these three residues in its ANTH domain. We note that in Figure 2C yap1802 3x PM-GFP shows an elevated signal at the bud neck, which is representative of the localization for this mutant. We speculate that since yap1802 3x PM-GFP is defective in both lipid and Snc2 binding, it is recruited to the bud neck by Ede1 through Yap1802’s NPF motifs because Ede1 is concentrated at the bud neck (Stimpson *et al*., 2009). Two-color imaging of the 3x PM and Ede1 confirmed strong colocalization between the two proteins at the bud neck (Figure S3C).

Since perturbing Yap1802’s interaction with lipids and cargo (Snc2) led to abnormal Yap1802 behavior, we investigated whether these endocytic perturbations also led to abnormal Snc2 behavior. Using confocal microscopy to visualize endogenously expressed mScarlet-Snc2 in yeast strains expressing point-mutated Yap1802-GFP fusion proteins, we observed differential distributions of Snc2 on the plasma membrane (Figure 2G). *yap1801Δ YAP1802* cells displayed normal Snc2 localization when compared to wild-type cells (compare Figure 2G with 1B), but each point mutant led to defects in Snc2 internalization, with an increase in Snc2 cortex signal in mother cells across all conditions (Figure 2H, Figure S3B). The Yap1802 K21E K23E mutant still shows a polarized distribution of Snc2 compared to other point mutants, but it also exhibits more plasma membrane signal in mothers compared to cells expressing wild-type Yap1802 (Figure 2G, H). This indicates that the Yap1802 K21E K23E mutant can still bind and internalize with Snc2 via CME, but the overall efficiency of this process is reduced, likely due to decreased lipid binding. This aligns with the reduced plasma membrane recruitment observed in the Yap1802 K21E K23E mutant (Figure 2C) and may lead to less Snc2 uptake via CME. Additionally, the largely plasma membrane-bound localization of Snc2 in the yap1802 L203S mutant further confirmed that the L203 residue of Yap1802 plays a key role in internalizing Snc2 through specific interactions (Figure 2G).

Taken together, these findings identify anionic phospholipids and Snc2 as important factors facilitating the recruitment of Yap1802 to the daughter plasma membrane. Since wild-type Yap1802 interacts with Snc2 directly and is most concentrated at the daughter cell tip where exocytosis is most active, Yap1802 localization is directly linked to polarized cell growth and secretion (Figure 3A). Signaling initiated by Cdc42 causes actin cable assembly from the daughter cell tip, enabling a type V myosin to recruit many cellular components, including secretory vesicles and the exocyst vesicle tethering complex to the bud tip (Figure 3A) (Pruyne and Bretscher, 2000a, 2000b; Mei and Guo, 2018; Mei *et al*., 2018). Once vesicles reach the plasma membrane, we suggest that Snc2 and anionic phospholipids act together as a “landing pad” for Yap1802, creating favorable conditions for its recruitment to the plasma membrane. Since Yap1802’s lipid binding and Snc2 binding display roughly equal importance for Yap1802 recruitment, we suggest that robust Yap1802 localization relies on its coincidence detection of these two factors (Figure 3B). Recent work has suggested that Yap1801/2 may be among the earliest-acting factors in polarized endocytosis, implicating this Yap1802/Snc2 interaction in the initiation of CME in daughter cells during cell growth (Sun *et al*., 2024). It seems likely that these properties would also be conserved for the highly homologous paralogs Yap1801 and Snc1, but this possibility remains to be tested.

**FIGURE 3:**
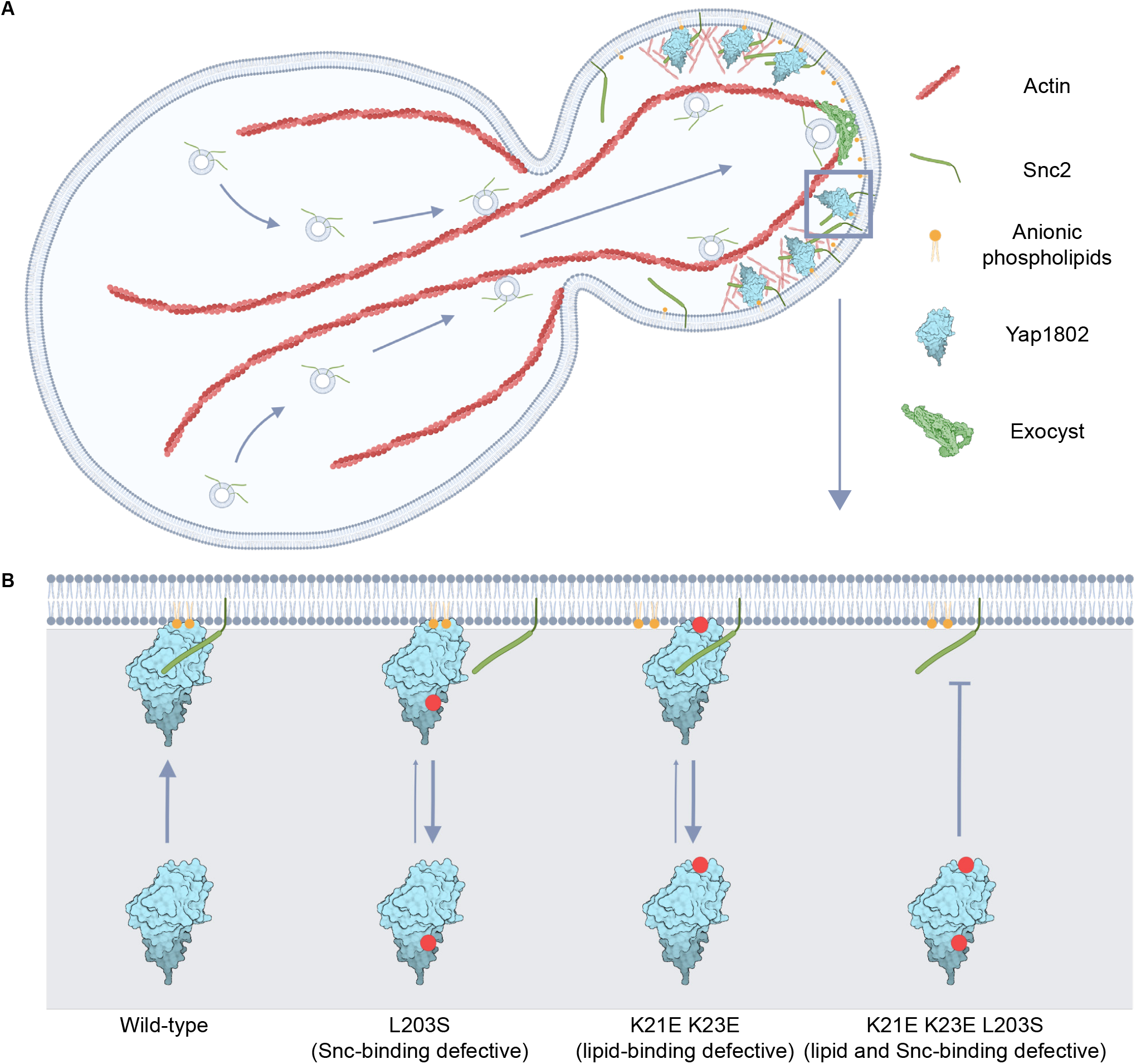
**A**. Model for how interaction between synaptobrevin-like v-SNARES Snc1/2 and the CME adaptor proteins Yap1801/2 contributes to CME site initiation and links endocytosis to polarized exocytosis. Cdc42-nucleated actin cables enable directional transport of vesicles toward the daughter bud tip, with fusion near the bud tip assisted by the exocyst vesicle tethering complex. This localized secretion enriches the daughter plasma membrane in Snc1/2, which, in concert with anionic phospholipids, recruit Yap1801/2 to internalize Snc1/2 via CME. **B**. Model for Snc1/2 and anionic phospholipid coincidence detection, enabling robust Yap1801/2 recruitment. Each individual Yap1802 mutant displays defects in recruitment when compared to wild-type, an effect most pronounced in the triple point mutant. Created with BioRender.com.

Interestingly, we could not precisely perturb the endocytic machinery in a way that led to cell growth defects like those observed in *snc2 V39A M42A/snc1Δ* cells. Since *yap1801Δ/yap1802Δ* double null cells also display essentially normal growth and viability, this result suggests the existence of adaptor proteins in addition to Yap1801/2 that can mediate Snc1/2 internalization, perhaps AP2 or Syp1 (Figure S2D) (Sun *et al*., 2024). In addition, the Hip1/R homolog Sla2, which also contains an ANTH domain, and the epsins Ent1/2, could be candidates for this role. However, since *sla2Δ* and *ent1Δ/ent2Δ* cells are completely defective in CME, and since our efforts to identify Sla2 or Ent1/2 residues that might mediate interactions between these proteins and Snc1/2 using Alphafold2 multimer analysis were not successful, studying the effect these proteins have on Snc1/2 internalization must await future studies and new approaches (Sun *et al*., 2005; Maldonado-Báez *et al*., 2008; Mirdita *et al*., 2022).

## Conclusions

Here, we provide evidence that the previously identified interactions between the endocytic adaptor Yap1802, the CME cargo Snc2, and anionic phospholipids provide a direct link between polarized cell growth driven by localized exocytosis and CME. The polarized delivery of Snc2 to the plasma membrane by exocytosis and the subsequent endocytosis of these proteins appear to be highly coordinated, as CME sites are concentrated near sites of localized exocytosis. Additionally, given the recent identification of Yap1801/2 as among the earliest-acting factors in CME, the Yap1802/Snc2 interaction could be involved in CME initiation. As the proteins investigated here are highly conserved in all eukaryotes, we expect that our observations and conclusions will prove relevant to other settings involving polarized secretion and efficient compensatory cargo retrieval via CME, such as at presynaptic nerve termini (Saheki and Camilli, 2012).

## Materials/Methods

### Strains, yeast husbandry, cloning, and plasmids

All yeast strains produced and analyzed in this study can be found in Table S1. All strains were derived from the wild-type diploid DDY1102 and were propagated using standard yeast husbandry techniques (Amberg *et al*., 2005). Genes encoding C-terminally-tagged fusion proteins were engineered as previously described (Longtine *et al*., 1998). Genes encoding N-terminally tagged Snc2 fusion proteins were engineered by cloning the genomic *SNC2* open reading frame (ORF) into a pFA6a-XFP backbone to create an in-frame fusion protein. PCR products with homology to the *SNC2* locus were generated from these plasmids, and then transformed into an *snc2Δ* background using previously described techniques (Longtine *et al*., 1998). The snc2 V39A M42A double mutant was made at the endogenous *SNC2* locus by introducing the appropriate mutations into the above plasmids using quick-change mutagenesis (Kunkel, 1985). Similarly, the yap1802 L203S mutant was made by introducing the appropriate mutation into the wild-type gene carried on a pFA6a-GFP backbone, then transforming the resulting PCR product into a *yap1802Δ* strain using previously described techniques (Longtine *et al*., 1998). All other point mutations were introduced using CRISPR-Cas9 (Anand *et al*., 2017). All plasmids can be found in Table S2.

### Yeast chemical fixation

Yeast strains were grown overnight to mid-log phase (roughly OD 0.2-0.4) in synthetic minimal media and fixed according to previous protocols in 4% paraformaldehyde solution in cytoskeleton buffer (Kaplan and Ewers, 2015).

### Live-cell imaging

Yeast strains were grown overnight to mid-log phase (roughly OD 0.2-0.4) in synthetic minimal media lacking tryptophan (imaging media, containing L-histidine, L-leucine, uracil, L-lysine, adenine, L-methionine, and 2% glucose), then adhered to 8-well imaging chambers (Cellvis C8-1.5H-N) or glass coverslips using 0.2 mg/ml Concanavalin A. *GFP-snc2 V39A M42A/snc1Δ* strains were grown overnight to mid-log phase in casamino acid media lacking tryptophan (0.17% yeast nitrogen base, 0.5% casamino acid, 0.5% ammonium sulfate, uracil, adenine, and 2% glucose) (Kurokawa *et al*., 2013). For CK-666 experiments, adhered cells were treated for 30 min. with 50 µM CK-666 in imaging media. Media was aspirated and washed once with imaging media before immediately commencing imaging. All imaging experiments were repeated a minimum of two times with biological replicates (one mat a and one mat alpha strain) with datapoints consisting either of individual cell intensity traces or individual endocytic events, as per figure legends.

Widefield microscopy was performed using a Nikon Eclipse Ti microscope equipped with a Nikon 100× 1.4 NA Plan Apo VC oil objective and a Neo 5.5 sCMOS camera from Andor Technology. GFP/mScarlet two-color imaging was performed using a Lumencore Spectra X LED light source with a Semrock FF-493/574-Di01 dual-pass dichroic and an FF01-524/628-25 dual-pass excitation filter. Nikon Elements software was used to control the system, and all imaging was done at room temperature (23°C).

Confocal imaging was performed using a Zeiss LSM 900 with Airyscan 2.0 detection equipped with a 60x 1.4 NA oil objective. 1x averaging was used for all acquisitions. Excitation laser wavelengths of 488 nm for GFP and 561 nm for mScarlet were used. Zeiss Zen Blue software was used to control the system, and imaging was performed at 25°C using a temperature-controlled chamber.

NSPARC confocal imaging was performed using a Nikon Eclipse Ti2 microscope equipped with a 60x 1.4 NA Plan Apo λD oil objective and an NSPARC (Nikon Spatial Array Confocal) detector. For the 488 nm laser, an emission range of 503-545 nm was used, and for the 561 nm laser, an emission range of 582-618 nm was used. All data were acquired in Galvo mode with a pinhole size of 18 µM.

### Immunoblotting

All protein samples were prepared from cells grown to mid-log phase using trichloroacetic acid (TCA) precipitation (Pedersen and Drubin, 2019). 10% SDS-polyacrylamide gels were run at 100 V and transferred to nitrocellulose membranes at 35 V for 12 hrs. Cells were blocked in 5% milk in PBS, primary antibodies were prepared in PBS with 0.05% milk and 0.1% Tween-20, and secondary antibodies were prepared in PBS with 0.005% milk and 0.1% Tween-20. All incubations occurred at RT for 1 hr with 3 x 5 min PBS washes in between. The following primary antibodies were used in this study: mouse anti-GFP 1:500 (RRID:AB_390913, 11814460001, Roche), mouse anti-V5 1:2,000 (RRID:AB_2792973, R960CUS, Thermo Fisher Scientific), rabbit anti-hexokinase 1:10,000 (RRID:AB_219918, 100-4159, Rockland), and mouse anti-PGK1 1:10,000 (RRID:AB_2532235, 459250, Thermo Fisher Scientific). Immune blots were probed with LICOR fluorophore-conjugated secondary antibodies and imaged on a LICOR Odyssey CLx.

### Purification of Yap1802 ANTH truncations

Expression and purification of wild-type ANTH (ANTH WT) and its K21E K23E mutant were carried out as follows. Plasmids encoding either ANTH WT or ANTH K21E K23E were transformed into *E. coli* BL21(DE3) competent cells. Following transformation, cells were cultured in 1 L of 2×YT medium at 37°C with shaking at 220 rpm until the optical density at 600 nm (OD_600_) reached ∼0.8. Protein expression was induced by adding 1 mM IPTG, and the cultures were incubated overnight at 18°C with shaking at 220 rpm. Cells were harvested by centrifugation at 4°C and resuspended in 40 mL of lysis buffer containing 25 mM HEPES (pH 7.4), 150 mM NaCl, 20 mM imidazole, 10 mM β-mercaptoethanol (β-ME), 1% Triton X-100, and one EDTA-free protease inhibitor tablet (Sigma-Aldrich). Cells were lysed by sonication on ice, and the lysates were clarified by centrifugation at 40,000 rpm for 40 minutes at 4°C. ANTH WT and K21E K23E proteins were found in the insoluble fraction. The pellets were resuspended in denaturing buffer (8 M urea, 25 mM HEPES pH 7.4, 150 mM NaCl, 20 mM imidazole) and centrifuged again under the same conditions. The denatured proteins, now in the supernatant, were incubated with Ni-NTA resin (G Biosciences, USA) for 1 hour at 4°C with gentle mixing. The resin was transferred to a gravity flow column and washed with the same urea-containing buffer. Bound proteins were eluted using elution buffer consisting of 25 mM HEPES, 150 mM NaCl, 250 mM imidazole, and 8 M urea (pH 7.4). Eluted proteins were gradually refolded by stepwise dialysis into buffers containing decreasing concentrations of urea (6 M, 4 M, 2 M, and finally 0 M) in 25 mM HEPES, 150 mM NaCl (pH 7.4) at 4°C overnight. The final protein products were aliquoted, flash-frozen in liquid nitrogen, and stored at -80°C.

### Purification of recombinant His_6_-HaloTag

His_6_-HaloTag was purified using the protocol in Ngo et al. 2025, with the following modifications. His_6_-HaloTag was cloned into a pET28a vector and expressed in *E. coli* Rosetta2(DE3)pLysS cells. A pre-culture (2.5 ml) was grown overnight at 37°C and diluted to a 250 ml culture. The culture was incubated at 37°C until the OD_600_ reached 0.6, and protein expression was induced upon addition of 50 µM IPTG for 4 hr at 37°C. The cells were harvested by centrifugation at 5,000xg for 10 min at 4°C, and the cell pellet was stored at - 80°C until use. All subsequent manipulations were performed at 4°C. The cell pellet was thawed on ice, resuspended in 7.5 ml of cold Ni-NTA lysis buffer (20 mM Tris-HCl pH 8.0, 300 mM NaCl, 10 mM imidazole, 2 mM MgCl_2_, and a protease inhibitor cocktail) and sonicated 5 times (5 sec on, 15 sec off, 20% power) using a Branson Sonifier 450. The lysate was clarified by centrifugation at 20,000xg for 20 min in a fixed angle FIBERlite F21-8×50y rotor, and the supernatant fraction was applied to a gravity flow column containing 0.5 ml of pre-equilibrated HisPur^™^ Ni-NTA Resin (Thermo Fisher Scientific). The column was washed with three column volumes of cold Ni-NTA wash buffer (similar recipe as Ni-NTA lysis buffer but with 25 mM imidazole) and eluted with 1.5 ml of cold Ni-NTA elution buffer (similar recipe as Ni-NTA lysis buffer but with 300 mM imidazole). The imidazole eluate was buffer exchanged into tris-buffered saline (TBS) using a HiTrap desalting column (Cytiva). The purified His_6_-HaloTag was distributed into aliquots, snap-frozen in liquid nitrogen, and stored at -80°C until use.

### Liposome preparation and sucrose density gradient flotation assays

Liposomes were prepared essentially as described in Williams et al., 2025, with the following modifications. Lipids (Avanti Polar Lipids) were mixed in a glass vial at the following ratios: 38% DOPC, 20% POPE, 20% DOPS, 3% PI(4,5)P_2_, 0.5% TexasRed-PE, and 18.5% cholesterol in 500 µl of 95% chloroform and 5% methanol. The lipid mixture was briefly placed in an incubator preheated to 60°C before being transferred to a vacuum chamber to evaporate the organic solvent. The dried lipids were hydrated in 500 µl of TBS via incubation at 60°C for 30 min with intermittent vortexing. The liposome suspension was then extruded through a polycarbonate filter of 200-nm pore size (Whatman). For sucrose density gradient flotation assays, binding reactions (60 µl) consisted of liposomes (20 µl), purified HaloTag (15-100 nM) or ANTH protein (see Table S3 for all Halo-Tag concentrations used and for computed concentrations of each ANTH mutant using densitometry), and TBS. The reactions were incubated at room temperature for 15 min, mixed with 60 µl of 60% sucrose in TBS (for a final concentration of 30% sucrose), and sequentially overlaid with 100 µl of 25% sucrose in TBS and 20 µl of TBS. The resulting sucrose step gradients were centrifuged in a TLA100 rotor (Beckman) at 100,000 rpm (436,000xg) for 10 min at 4°C. The floated liposomes were collected, and the amount of fluorescent lipid recovered was quantified using a Tecan Infinite M Plex plate reader (Tecan). The bound proteins were analyzed by SDS-PAGE and immunoblotting after adjusting the sample volume to normalize for lipid recovery. Antibodies used for immunoblotting were mouse anti-6*His (RRID:AB_2923721, 66005-1-Ig, Proteintech) with sheep anti-mouse secondary (RRID:AB_772209, NXA931, Cytiva) and mouse anti-His-Tag HRP conjugate (RRID:AB_2797714, 9991, Cell Signaling Technology). Blots were developed with chemiluminescence and imaged on a BioRad ChemiDoc. Replicates from both primary antibodies were combined for relative protein binding measurements, and replicates using only the Proteintech antibody were used for concentration calibration and binding fraction measurements.

### Image and data analysis (code availability)

Montages and kymographs were generated using custom scripts in Fiji (NIH) as well as the open-source cell segmentation package CellPose (Stringer *et al*., 2021; Sun *et al*., 2024, 2017; Lu and Drubin, 2017). Line trace intensity data for Yap1802 point mutants was analyzed using custom MATLAB scripts. Line traces of medium-budded cells were generated using CellPose, then each intensity profile was interpolated to match the length of the mean trace length. Intensity profiles were averaged at each length and plotted using the MATLAB package plot_areaerrorbar (Martínez-Cagigal, 2024). Moving arrowheads in supplemental movies were added using the Draw_Arrows plugin (Daetwyler *et al*., 2020). Plots were generated with MATLAB and GraphPad Prism 10. All code is available at https://github.com/DrubinBarnes/Marchando_2024_Snc and https://github.com/geanhu/cme-movie-analysis-2024.

## Acknowledgements

We thank all members of the Drubin/Barnes laboratory for their help and support. We thank Randy Schekman for allowing one of us (JMN) to perform lipid binding experiments for this study. Yuichiro Iwamoto and Ross Pedersen provided constructive feedback and careful reading of our manuscript. We thank Andrew Wong for his assistance in generating point mutant strains. We are grateful to Constantine Bartolutti and Cyrus Ruediger of the Elçin Ünal/Gloria Brar laboratory for their assistance with CRISPR/Cas9 methodology and for sharing the necessary reagents. In addition, thank you to James Mester for conducting the Nikon NSPARC imaging experiments. This work was supported by NIH grant R35GM118149 (D.G. Drubin).

## Figure Legends

**FIGURE S1:**
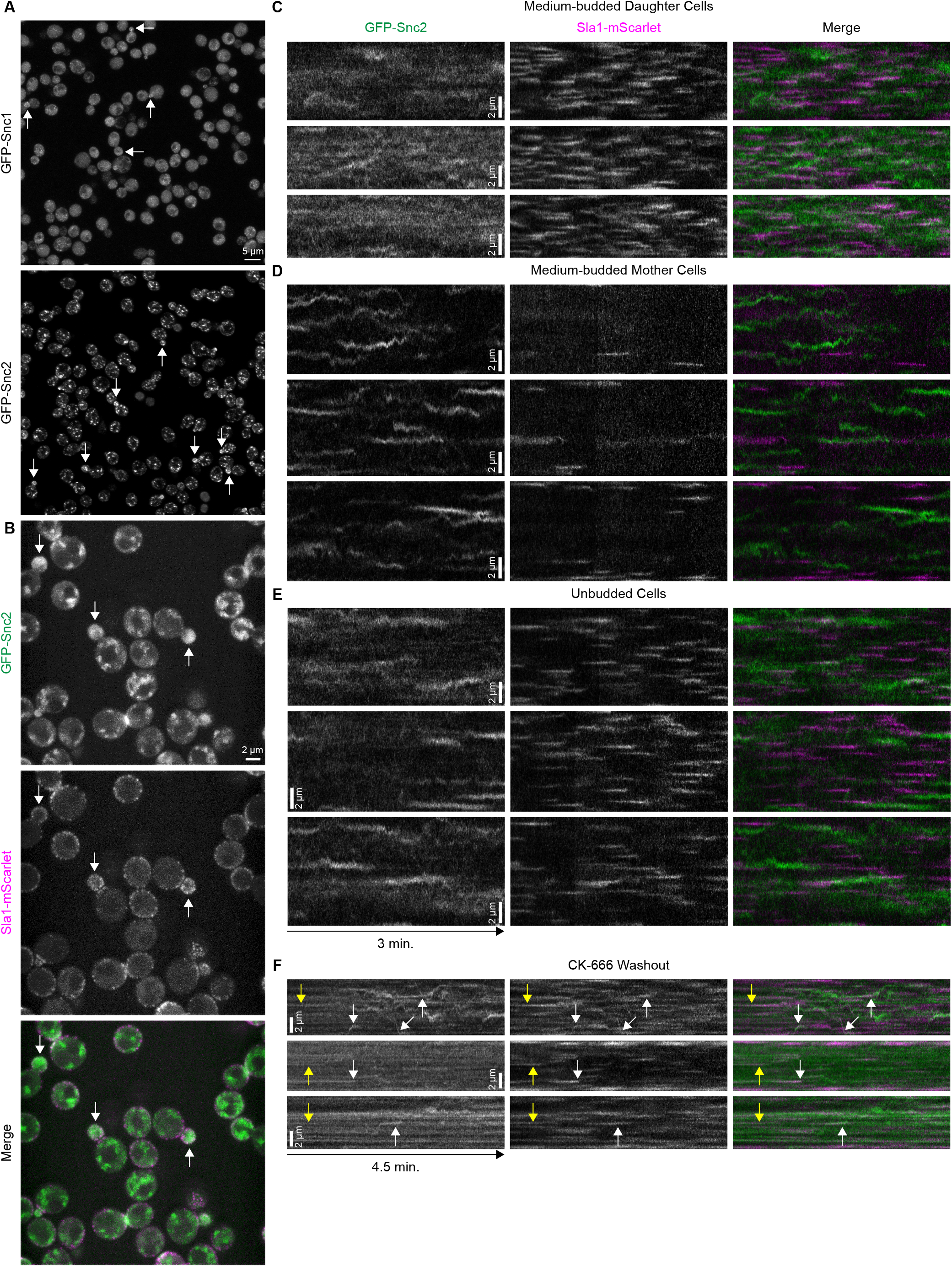
**A**. Maximum intensity projections of Airyscan confocal z-stacks of fixed yeast cells expressing GFP-Snc1 (with intron removed from the *SNC1* gene) and GFP-Snc2, respectively. Budded cells are demarcated with arrows. **B**. Maximum intensity projections from a time-lapse NSPARC movie of cells expressing GFP-Snc2 and Sla1-mScarlet. Budded cells are demarcated with arrows. **C**., **D**., **E**. Three examples each of circumferential kymographs made from 3-minute NSPARC movies of cells expressing GFP-Snc2 and Sla1-mScarlet in daughter, mother, and unbudded cells, respectively. **F**. Circumferential kymographs made from a 4.5-minute NSPARC movie of daughter cells expressing GFP-Snc2 and Sla1-mScarlet acquired directly after washout of 50 µM CK-666 following 30 min. of treatment. White and yellow arrows denote distinct Snc2/Sla1 interaction events.

**FIGURE S2:**
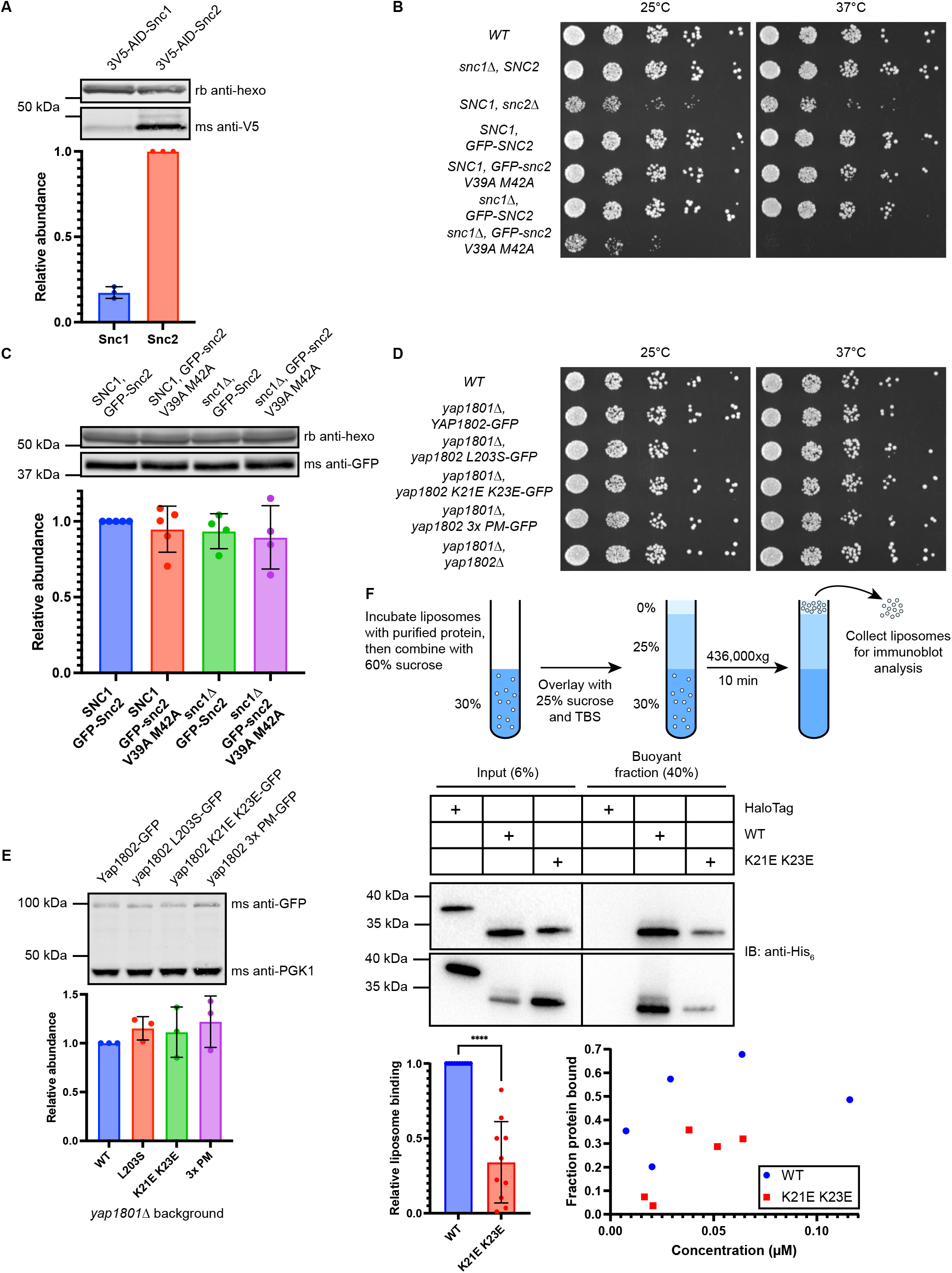
**A**. Immunoblot displaying relative levels of V5-AID-tagged Snc1 and Snc2 expressed from their normal promoters at their endogenous loci with anti-hexokinase loading control. **B**. Serial dilutions of strains carrying indicated *snc* null and point mutants and fluorescent fusion proteins on YPD. **C**. Immunoblot displaying relative levels of GFP-tagged Snc2 and snc2 V39A M42A mutant in *SNC1* and *snc1Δ* backgrounds. **D**. Serial dilutions of strains carrying various Yap1802 point mutations in a *yap1801Δ* background on YPD. **E**. Immunoblot displaying relative levels of GFP-tagged Yap1802 point mutants. **F**. Liposome flotation assays with purified truncated Yap1802 and yap1802 K21E K23E ANTH domains, two different replicates with different protein amounts loaded. Liposomes were made with a molar composition of 38% DOPC, 20% POPE, 20% DOPS, 3% PI(4,5)P_2_, 0.5% TexasRed-PE, and 18.5% cholesterol. Relative binding was computed for each replicate with one exposure optimized for the input and one optimized for the buoyant fraction. Effective concentrations and binding fractions of the two ANTH proteins were determined using densitometry in relation to the known His_6_-HaloTag concentration in each replicate using a single optimized exposure. Error bars for all immunoblots represent standard deviation. **** indicates p < 0.0001.

**MOVIES S1 and S2:** Snc2 interacts with the endocytic machinery in a complex manner. Three-minute movies acquired using a Nikon confocal microscope equipped with an NSPARC detector (see Materials and Methods) of live cells expressing GFP-Snc2 (green) and Sla1-mScarlet (magenta), both from the endogenous promoter and locus from the endogenous promoter and locus in an *snc1Δ* background. Moving arrowheads track distinct Snc2-positive compartments. **Movie S1:** first, a compartment translocating from one location on the plasma membrane to another in a daughter cell, and second, two compartments in the mother cell that move directionally from the cytosol to the bud neck. **Movie S2:** Moving arrowheads track distinct interaction events between GFP-Snc2 and Sla1-mScarlet: first and second at the daughter cell cortex, and third in a mother cell. Acquisition temporal resolution is ∼0.6 seconds, playback rate is 27 frames per second. Scale bar is 2 µm.

**FIGURE S3:**
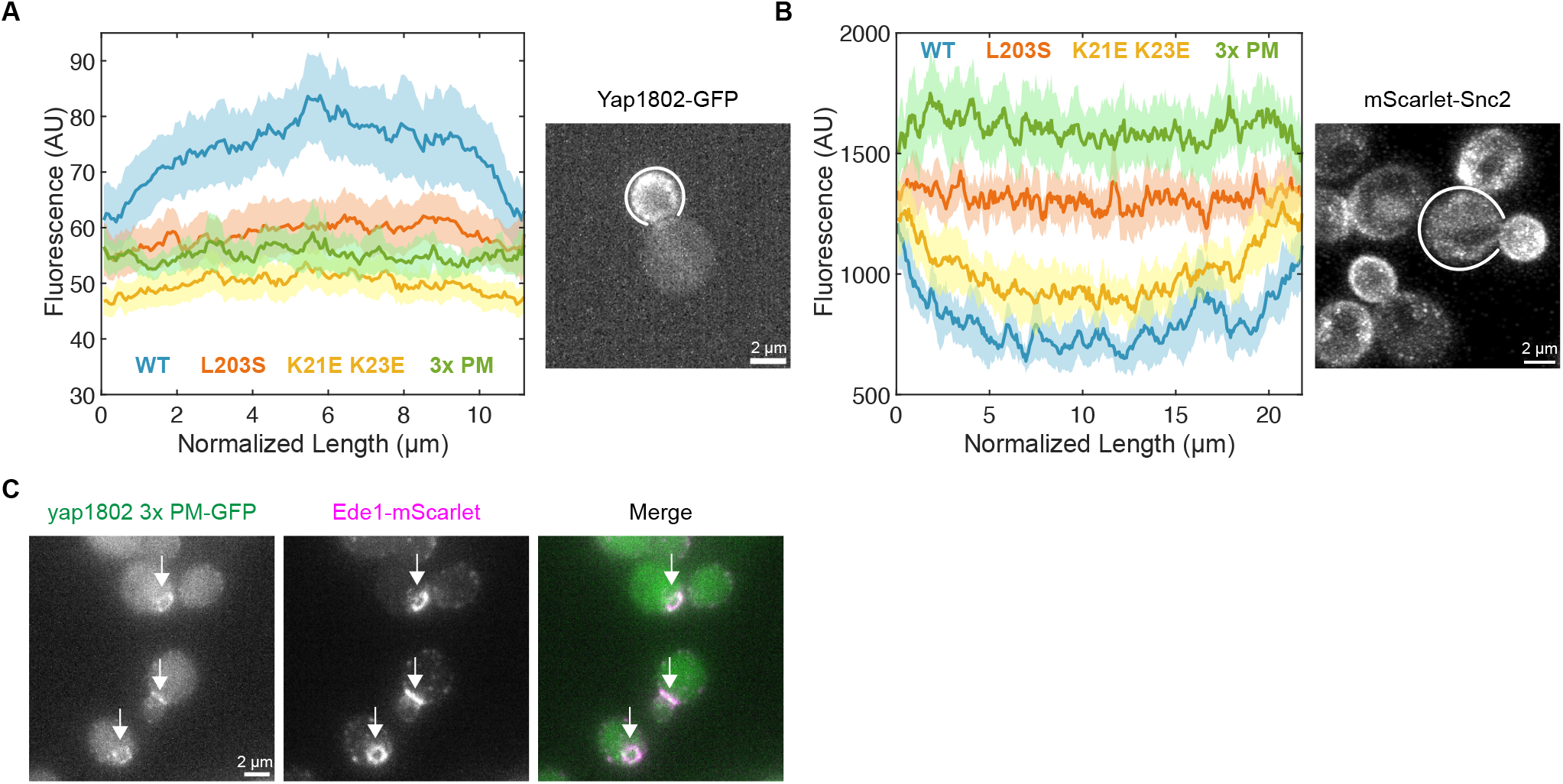
**A**. GFP line traces of daughter cells as in **C** from wild-type and Yap1802 point mutant maximum intensity projections for the same strains used in **Figure 2C. B**. mScarlet-Snc2 line traces from maximum intensity projections of mother cells as in **Figure 2G** expressing Yap1802 wild-type protein or point mutants. For each plot, each trace from individual cells was interpolated to match the mean length of cells in the wild-type cohort. Outlines represent a 95% confidence interval for each point in the trace, annotated microscopy images display examples of individual line traces used. **C**. Maximum intensity projections of widefield z-stacks of live cells expressing yap1802 3x PM-GFP and Ede1-mScarlet in a *yap1801Δ* background. Colocalization is denoted with arrows.

