## Supplemental Tables 1, 2, and 3 for "Polarized anionic phospholipids and exocytosis are implicated in the polarized recruitment of budding yeast AP180, an endocytic initiator"

| DDY Number | Alias | Mat | Base Genotype | Additional Genotype | Background |
|---|---|---|---|---|---|
| 5943 | PMY314.1 | A | his3Δ200, ura3-52, leu2-3, 112, lys2-801(am)? | *GFP-SNC1 (no intron)::cgLEU2* | DDY1102 |
| 5944 | PMY288.1 | A | his3Δ200, ura3-52, leu2-3, 112, lys2-801(am)? | *GFP-SNC2::cgLEU2* | DDY1102 |
| 5945 | PMY317.1 | A | his3Δ200, ura3-52, leu2-3, 112, lys2-801(am)? | *3V5-AID2-SNC1 (no intron)::KanMX, TIR1::LEU2* | DDY1102 |
| 5946 | PMY292.2 | A | his3Δ200, ura3-52, leu2-3, 112, lys2-801(am)? | *3V5-AID2-SNC2::KanMX, TIR1::LEU2* | DDY1102 |
| 5947 | PMY277.1 | A | his3Δ200, ura3-52, leu2-3, 112, lys2-801(am)? | *snc1Δ::cgHIS3* | DDY1102 |
| 5948 | PMY281.1 | A | his3Δ200, ura3-52, leu2-3, 112, lys2-801(am)? | *snc2Δ::cgHIS3* | DDY1102 |
| 5949 | PMY311.1 | A | his3Δ200, ura3-52, leu2-3, 112, lys2-801(am)? | *GFP-SNC2::cgLEU2, snc1Δ::cgHIS3, SLA1-yomScarlet-I::KanMX* | DDY1102 |
| 5950 | PMY312.1 | α | his3Δ200, ura3-52, leu2-3, 112, lys2-801(am)? | *GFP-SNC2::cgLEU2, snc1Δ::cgHIS3, SLA1-yomScarlet-I::KanMX* | DDY1102 |
| 5951 | PMY290.1 | A | his3Δ200, ura3-52, leu2-3, 112, lys2-801(am)? | *GFP-SNC2 V39A M42A::cgLEU2* | DDY1102 |
| 5952 | PMY409.1 | A | his3Δ200, ura3-52, leu2-3, 112, lys2-801(am)? | *GFP-SNC2 V39A M42A::cgLEU2, YAP1802-yomScarlet-I::KanMX, snc1Δ::cgHIS3* | DDY1102 |
| 5953 | PMY374.1 | α | his3Δ200, ura3-52, leu2-3, 112, lys2-801(am)? | *YAP1802 L203S-GFP::cgLEU2, SLA1-yomScarlet-I::KanMX, yap1801Δ::NatR* | DDY1102 |
| 5954 | PMY375.1 | A | his3Δ200, ura3-52, leu2-3, 112, lys2-801(am)? | *YAP1802 L203S-GFP::cgLEU2, SLA1-yomScarlet-I::KanMX, yap1801Δ::NatR* | DDY1102 |
| 5955 | PMY376.1 | α | his3Δ200, ura3-52, leu2-3, 112, lys2-801(am)? | *YAP1802 K21E K23E-GFP::HIS3, SLA1-yomScarlet-I::KanMX, yap1801Δ::NatR* | DDY1102 |
| 5956 | PMY377.1 | A | his3Δ200, ura3-52, leu2-3, 112, lys2-801(am)? | *YAP1802 K21E K23E-GFP::HIS3, SLA1-yomScarlet-I::KanMX, yap1801Δ::NatR* | DDY1102 |
| 5957 | PMY378.1 | α | his3Δ200, ura3-52, leu2-3, 112, lys2-801(am)? | *YAP1802 K21E K23E L203S-GFP::cgLEU2, SLA1-yomScarlet-I::KanMX, yap1801Δ::NatR* | DDY1102 |
| 5958 | PMY379.1 | A | his3Δ200, ura3-52, leu2-3, 112, lys2-801(am)? | *YAP1802 K21E K23E L203S-GFP::cgLEU2, SLA1-yomScarlet-I::KanMX, yap1801Δ::NatR* | DDY1102 |
| 5959 | PMY382.1 | α | his3Δ200, ura3-52, leu2-3, 112, lys2-801(am)? | *YAP1802-GFP::HIS3, SLA1-yomScarlet-I::KanMX, yap1801Δ::NatR* | DDY1102 |
| 5960 | PMY383.1 | A | his3Δ200, ura3-52, leu2-3, 112, lys2-801(am)? | *YAP1802-GFP::HIS3, SLA1-yomScarlet-I::KanMX, yap1801Δ::NatR* | DDY1102 |
| 5961 | PMY423.1 | A | his3Δ200, ura3-52, leu2-3, 112, lys2-801(am)? | *YAP1802 K21E K23E L203S-GFP::cgLEU2, yomScarlet-I-SNC2::cgURA3, yap1801Δ::NatR* | DDY1102 |
| 5962 | PMY441.1 | A | his3Δ200, ura3-52, leu2-3, 112, lys2-801(am)? | *YAP1802-GFP::HIS3, yomScarlet-I-SNC2::cgURA3, yap1801Δ::NatR* | DDY1102 |
| 5963 | PMY451.1 | A | his3Δ200, ura3-52, leu2-3, 112, lys2-801(am)? | *YAP1802 L203S-GFP::cgLEU2, yomScarlet-I-SNC2::cgURA3, yap1801Δ::NatR* | DDY1102 |
| 5964 | PMY452.1 | α | his3Δ200, ura3-52, leu2-3, 112, lys2-801(am)? | *YAP1802 K21E K23E-GFP::HIS3, yomScarlet-I-SNC2::cgURA3, yap1801Δ::NatR* | DDY1102 |
| 5971 | PMY462.1 | α | his3Δ200, ura3-52, leu2-3, 112, lys2-801(am)? | *YAP1802 K21E K23E L203S-GFP::cgLEU2, EDE1-yomScarlet-I::KanMX, yap1801Δ::NatR* | DDY1102 |
| 5972 | PMY463.1 | A | his3Δ200, ura3-52, leu2-3, 112, lys2-801(am)? | *YAP1802 K21E K23E L203S-GFP::cgLEU2, EDE1-yomScarlet-I::KanMX, yap1801Δ::NatR* | DDY1102 |

| pDD number | Name | Backbone | Resistance | Contents | Purpose | Competent Cells |
|---|---|---|---|---|---|---|
| pDD2745 | pPM129 | pFA6a | Amp | GFP-SNC2::cgLEU2 | Longtine-style tagging plasmid | DH5α |
| pDD2746 | pPM131 | pFA6a | Amp | GFP-SNC2 V39A M42A::cgLEU2 | Longtine-style tagging plasmid | DH5α |
| pDD2747 | pPM133 | pFA6a | Amp | 3V5-AID2-SNC2::KanMX | Longtine-style tagging plasmid | DH5α |
| pDD2748 | pPM134 | pFA6a | Amp | GFP-SNC1 (no intron)::cgLEU2 | Longtine-style tagging plasmid | DH5α |
| pDD2749 | pPM135 | pFA6a | Amp | 3V5-AID2-SNC1 (no intron)::KanMX | Longtine-style tagging plasmid | DH5α |
| pDD2750 | pPM150 | pFA6a | Amp | YAP1802 L203S-GFP::cgLEU2 | Longtine-style tagging plasmid | DH5α |
| pDD2751 | pPM152 | pUB1306 | Kan | CAS9-GTCTTCATTATATACTAAGT (YAP1802 gRNA)::URA3 CEN | Yeast expression plasmid for YAP1802 editing mediated by CAS9 | DH5α |
| pDD2752 | pPM163 | pFA6a | Amp | yomScarlet-I-SNC2::cgURA3 | Longtine-style tagging plasmid | DH5α |
| pDD2753 | pPM164 | MacroLab 1-B | Kan | YAP1802 ANTH-6His | Expression vector for purification | DH5α |
| pDD2754 | pPM165 | MacroLab 1-B | Kan | YAP1802 ANTH K21E K23E-6His | Expression vector for purification | DH5α |

| Replicate | HaloTag amount (18 µM stock) | WT amount | 2K-E amount | Date |
|---|---|---|---|---|
| 1 | 2 | 5 | 5 | 8/6/25 |
| 1:5 dilution | | 5 | 5 | |
| | | 5 | 5 | |
| | | 5 | 5 | |
| 2 | 2 | 5 | 2.2 | 8/7/25 |
| 1:15 dilution | | 5 | 2.2 | |
| | | 5 | 2.2 | |
| | | 5 | 2.2 | |
| 3 | 2 | 5 | 2.2 | 8/7/25 |
| 1:15 dilution | | 5 | 2.2 | |
| | | 5 | 2.2 | |
| | | 5 | 2.2 | |
| 4 | 2 | 3 | 2.5 | 8/21/25 |
| 1:15 dilution | | 3 | 2.5 | |
| | | 3 | 2.5 | |
| | | 3 | 2.5 | |
| 5 | 2 | 3 | 2.5 | 8/24/25 |
| None | | 3 | 2.5 | |
| | | 3 | 2.5 | |
| | | 3 | 2.5 | |
| | | 3 | 2.5 | |
| | | 3 | 2.5 | |
| 6 | 2 | 3 | 2.5 | 10/9/25 |
| 1:15 dilution | | 3 | 2.5 | |
| | | 3 | 2.5 | |
| | | 3 | 2.5 | |
| 7 | 2 | 3 | 2.5 | 10/9/25 |
| 1:15 dilution | | 3 | 2.5 | |
| | | 3 | 2.5 | |
| | | 3 | 2.5 | |
| 8 | 2 | 3 | 2.5 | 10/10/25 |
| 1:5 dilution | | 3 | 2.5 | |
| | | 3 | 2.5 | |
| | | 3 | 2.5 | |
| | | 3 | 2.5 | |
| | | 3 | 2.5 | |
| 9 | 2 | 3 | 3 | 10/28/25 |
| 1:5 dilution | | 3 | 3 | |
| | | 3 | 3 | |
| | | 3 | 3 | |
| | | 3 | 3 | |
| | | 3 | 3 | |
| | | 3 | 3 | |
| | | 3 | 3 | |
| 10 | 2 | 3 | 2 | 10/31/25 |
| 1:15 dilution | | 3 | 2 | |
| | | 3 | 2 | |
| | | 3 | 2 | |
| 11 | 2 | 3 | 3 | 1/28/26 |
| 1:15 dilution | | 3 | 3 | |
| 12 | 1 | 4.5 | 2 | 1/29/26 |
| 1:15 dilution | | 4.5 | 2 | |

This means saturated signal

**6His-Halo MW**

**ANTH MW**
**2K-E MW**

| Analysis type | Exposure time (sec) | Halo input intensity | WT input intensity | 2K-E input intensity |
|---|---|---|---|---|
| Volume analysis | 76.2 | 22,195,224.90 | 7,044,512.34 | 12,524,787.03 |
| | 34.2 | 12,441,270.14 | 3,139,094.87 | 5,589,927.89 |
| Band analysis | 76.2 | 34759050 | 11657300 | 17973500 |
| | 34.2 | 17319350 | 4265100 | 7278300 |
| Volume analysis | 35.2 (high sensitivity) | 2,158,933.30 | 3,438,688.93 | 2,167,459.35 |
| | 35.2 | 2,084,998.29 | 3,457,105.33 | 2,065,669.14 |
| Band analysis | 35.2 (high sensitivity) | 2622115 | 4831817 | 2734838 |
| | 35.2 | 2325100 | 4133800 | 2411500 |
| Volume analysis | 67.3 | 5,980,969.11 | 17,401,457.47 | 9,964,611.49 |
| | 21.1 | 1,829,342.38 | 5,518,325.19 | 3,057,723.57 |
| Band analysis | 67.3 | 6229548 | 25414065 | 13337163 |
| | 21.1 | 1792648 | 6542796 | 3644628 |
| Volume analysis | 63 | 680,233.45 | 2,519,811.64 | 4,150,836.73 |
| | 15 | 158,063.27 | 622,733.45 | 974,920.00 |
| Band analysis | 63 | 833846 | 3250728 | 5319678 |
| | 15 | 200954 | 678002 | 1150292 |
| Volume analysis | 299.98 | 13,152,363.63 | 2,509,271.50 | 1,199,448.38 |
| | 91.6 | 9,166,460.63 | 729,544.63 | 369,765.38 |
| | 31.2 | 4,166,565.88 | 215,723.88 | 105,272.13 |
| Band analysis | 299.98 | 22008570 | 2817638 | 1339658 |
| | 91.6 | 10720620 | 945162 | 405,513.00 |
| | 31.2 | 4784850 | 283659 | 132572 |
| Volume analysis | 97 | 878,556.62 | 13,001,218.81 | 7,943,284.94 |
| | 26 | 179,766.03 | 3,358,658.91 | 2,077,268.83 |
| Band analysis | 97 | 766,536.00 | 15,311,644.00 | 9,248,212.00 |
| | 26 | 145,464.00 | 3,628,430.00 | 2,096,346.00 |
| Volume analysis | 29 | 187,259.29 | 3,668,332.57 | 1,563,737.86 |
| | 12 | 21,261.56 | 1,413,553.56 | 612,130.44 |
| Band analysis | 29 | 224884 | 4466066 | 1897104 |
| | 12 | 51198 | 1549674 | 583926 |
| Volume analysis | 200 | 7,348,355.80 | 2,428,143.20 | 2,335,569.60 |
| | 43.2 | 2,898,624.00 | 502,300.00 | 485,472.00 |
| | 21.1 | 1,428,089.60 | 253,252.00 | 233,506.40 |
| Band analysis | 200 | 10,895,871.00 | 2,909,571.00 | 2,808,183.00 |
| | 43.2 | 3,578,547.00 | 557,718.00 | 548,289.00 |
| | 21.1 | 1726074 | 282618 | 261744 |
| Volume analysis | 654.76 | 7,795,392.71 | 195,216.57 | 849,265.14 |
| | 391.4 | 8,636,647.14 | 177,788.57 | 641,512.57 |
| | 119 | 4,217,156.43 | 37,817.71 | 144,012.00 |
| | 55.5 | 2,284,928.00 | 23,084.00 | 57,585.71 |
| Band analysis | 654.76 | 9,318,188.00 | 671,220.00 | 932,712.00 |
| | 391.4 | 10,501,172.00 | 467896 | 692780 |
| | 119 | 4534772 | 110,902.00 | 211,552.00 |

| | | | | |
|---|---|---|---|---|
| | 55.5 | 2,438,656.00 | 56,452.00 | 103,554.00 |
| Volume analysis | 600 | 905,447.61 | 391,833.55 | -81,442.61 |
| | 13.1 | 92,673.29 | 54,446.45 | 35,227.87 |
| Band analysis | 600 | 1288854 | 445473 | 317100 |
| | 13.1 | 136101 | 85071 | 42840 |
| Volume analysis | 7 | 18,454,558.34 | 3,637,687.24 | 9,908,783.73 |
| Band analysis | 7 | 23,071,160.00 | 4,802,784.00 | 13680212 |
| Volume analysis | 22.1 | 11,628,035.34 | 12,214,961.97 | 9,967,034.03 |
| Band analysis | 22.1 | 14,577,097.00 | 21,698,725.00 | 13,739,745.00 |

37

33
33

| WT flotation intensity | 2K-E flotation intensity | 2K-E/WT float ratio | WT/Halo ratio |
|---|---|---|---|
| 21,331,014.66 | 21,696,355.10 | 1.017127194 | 0.317388644 |
| 12,594,807.04 | 11,239,288.85 | 0.892374834 | 0.252313055 |
| 40682650 | 39255250 | 0.96491379 | 0.335374528 |
| 21763600 | 18102650 | 0.831785642 | 0.246262129 |
| 8,344,392.33 | 5,013,533.91 | 0.600826724 | 1.592772193 |
| 16,391,762.67 | 5,172,084.76 | 0.315529505 | 1.658085456 |
| 16753716 | 7063599 | 0.421613868 | 1.842717425 |
| 22655800 | 7071900 | 0.312145234 | 1.777902026 |
| 35,772,838.77 | 22,976,959.36 | 0.642301818 | 2.909471218 |
| 18,753,612.60 | 6,840,666.72 | 0.364765279 | 3.016562259 |
| 52963398 | 29159964 | 0.550568224 | 4.079600157 |
| 23979384 | 9969024 | 0.415733115 | 3.649794048 |
| 11,160,774.82 | 6,850,345.73 | 0.613787648 | 3.704333592 |
| 4,730,561.82 | 1,584,198.91 | 0.334885997 | 3.939773255 |
| 18651390 | 10060960 | 0.539421459 | 3.898475258 |
| 5841576 | 2106416 | 0.360590361 | 3.373916419 |
| 11,063,175.50 | 2,003,539.13 | 0.181099823 | 0.190784833 |
| 4,587,951.13 | 542,215.50 | 0.118182493 | 0.079588475 |
| 1,533,209.88 | 173,092.25 | 0.11289534 | 0.051774982 |
| 16644376 | 2927745 | 0.175899956 | 0.128024583 |
| 5,449,836.00 | 810,999.00 | 0.148811634 | 0.088162998 |
| 1730820 | 285066 | 0.164699969 | 0.059282736 |
| 31,675,399.70 | 25,857,259.14 | 0.816319901 | 14.7983846 |
| 15,340,140.77 | 7,718,990.70 | 0.50318904 | 18.68350202 |
| 48,141,630.00 | 33,166,742.00 | 0.688940985 | 19.97511402 |
| 19197106 | 9501140 | 0.494925641 | 24.9438349 |
| 10,760,460.29 | -370,924.43 | -0.034471056 | 19.58958968 |
| 15,883,864.39 | -199,510.93 | -0.012560604 | 66.48399723 |
| 18224472 | 1119932 | 0.061452096 | 19.85942086 |
| 17861298 | 122598 | 0.006863891 | 30.26825267 |
| 5,548,226.80 | 689,504.60 | 0.124274768 | 0.330433537 |
| 6,695,349.40 | 65,992.80 | 0.009856513 | 0.173289119 |
| 3,678,841.60 | 35,845.60 | 0.009743719 | 0.177336212 |
| 13,586,748.00 | 1,676,850.00 | 0.123418054 | 0.267034274 |
| 7,947,786.00 | 292551 | 0.036809119 | 0.155850405 |
| 4275999 | 153,468.00 | 0.03589056 | 0.163734579 |
| 8,713,548.71 | 2,387,030.43 | 0.273944693 | 0.025042558 |
| 9,145,666.43 | 2,374,810.29 | 0.259665089 | 0.020585369 |
| 3,865,338.29 | 495,473.14 | 0.128183643 | 0.008967586 |
| 1,848,749.14 | 247,335.43 | 0.133785284 | 0.010102725 |
| 9,935,992.00 | 3330910 | 0.335236784 | 0.072033318 |
| 9598820 | 3089438 | 0.32185602 | 0.04455655 |
| 4,228,840.00 | 767,866.00 | 0.1815784 | 0.024455915 |

| | | | |
|---|---|---|---|
| 1,857,724.00 | 330968 | 0.178157789 | 0.023148816 |
| 3,603,396.10 | 2,446,877.29 | 0.67904755 | 0.432751208 |
| 1,947,184.39 | 274,965.29 | 0.141211737 | 0.587509642 |
| 13651995 | 4282299 | 0.313675694 | 0.345634959 |
| 3009384 | 475986 | 0.158167253 | 0.625057861 |
| 8,999,684.13 | 2,529,661.80 | 0.281083398 | 0.19711592 |
| 11908519 | 2877770 | 0.241656414 | 0.208172628 |
| 17,199,755.34 | 5,155,444.46 | 0.299739406 | 1.050475133 |
| 26,232,204.00 | 6,595,416.00 | 0.251424394 | 1.488549126 |

| 2K-E/Halo ratio | HaloTag concentration (µM) | WT concentration back-calculated (µM) |
|---|---|---|
| 0.564300974 | 0.102857143 | 0.077173725 |
| 0.449305242 | | 0.077173725 |
| 0.517088355 | | 0.077173725 |
| 0.420240944 | | 0.077173725 |
| 1.003949194 | 0.034285714 | 0.077173725 |
| 0.99072942 | | 0.077173725 |
| 1.042989343 | | 0.077173725 |
| 1.037159692 | | 0.077173725 |
| 1.666052994 | 0.034285714 | 0.077173725 |
| 1.671487854 | | 0.077173725 |
| 2.14095196 | | 0.077173725 |
| 2.033097407 | | 0.077173725 |
| 6.10207672 | 0.034285714 | 0.046304235 |
| 6.167909744 | | 0.046304235 |
| 6.379688815 | | 0.046304235 |
| 5.724155777 | | 0.046304235 |
| 0.091196412 | 0.514285714 | 0.046304235 |
| 0.040338948 | | 0.046304235 |
| 0.025265921 | | 0.046304235 |
| 0.060869834 | | 0.046304235 |
| 0.037825518 | | 0.046304235 |
| 0.027706616 | | 0.046304235 |
| 9.041289683 | 0.034285714 | 0.046304235 |
| 11.55540275 | | 0.046304235 |
| 12.0649415 | | 0.046304235 |
| 14.41144201 | | 0.046304235 |
| 8.35065589 | 0.034285714 | 0.046304235 |
| 28.79047497 | | 0.046304235 |
| 8.43592252 | | 0.046304235 |
| 11.40525021 | | 0.046304235 |
| 0.317835671 | 0.102857143 | 0.046304235 |
| 0.167483606 | | 0.046304235 |
| 0.163509629 | | 0.046304235 |
| 0.257729098 | | 0.046304235 |
| 0.153215537 | | 0.046304235 |
| 0.151641239 | | 0.046304235 |
| 0.108944498 | 0.102857143 | 0.046304235 |
| 0.074277965 | | 0.046304235 |
| 0.034149077 | | 0.046304235 |
| 0.02520242 | | 0.046304235 |
| 0.100095856 | | 0.046304235 |
| 0.065971684 | | 0.046304235 |
| 0.046651077 | | 0.046304235 |

| | | |
|---|---|---|
| 0.042463554 | | 0.046304235 |
| -0.08994735 | 0.034285714 | 0.046304235 |
| 0.38012971 | | 0.046304235 |
| 0.246032522 | | 0.046304235 |
| 0.31476624 | | 0.046304235 |
| 0.536928793 | 0.034285714 | 0.046304235 |
| 0.592957268 | | 0.046304235 |
| 0.857155464 | 0.017142857 | 0.069456352 |
| 0.942557013 | | 0.069456352 |

| WT concentration (μM) | WT binding percentage (same blot) | Relative Binding |
|---|---|---|
| 0.036602742 | 7.570082086 | |
| 0.029097921 | 10.030604 | 0.501912369 |
| 0.038676959 | 8.724715414 | |
| 0.0284001 | 12.75679351 | 0.539481724 |
| 0.061228645 | 6.066550722 | |
| 0.063739389 | 11.85367604 | 0.500589696 |
| 0.07083693 | 8.66843467 | |
| 0.068345325 | 13.70155789 | 0.551487382 |
| 0.111844608 | 5.1393452 | |
| 0.115961354 | 8.496061733 | 0.636998993 |
| 0.156826188 | 5.210047861 | |
| 0.140303771 | 9.162514008 | 0.792182595 |
| 0.142400356 | 11.07302492 | |
| 0.151451024 | 18.99111804 | 0.203296272 |
| 0.149863464 | 14.34401002 | |
| 0.129698605 | 21.53967097 | 0.220348146 |
| 0.110010995 | 11.0222982 | |
| 0.045892575 | 15.72196877 | 0.247240287 |
| 0.029854665 | 17.76819876 | 0.222741755 |
| 0.073821967 | 14.76802201 | |
| 0.050836846 | 14.4150844 | 0.312988326 |
| 0.034183811 | 15.25440758 | 0.38387956 |
| 0.568872966 | 6.090851976 | |
| 0.718222935 | 11.41835267 | 0.82359765 |
| 0.767874513 | 7.860297366 | |
| 0.958879887 | 13.22686809 | 0.819415172 |
| 0.753054357 | 7.333345652 | |
| 2.555748465 | 28.09208089 | |
| 0.763427087 | 10.20163607 | |
| 1.163558804 | 28.81460552 | 0.016158624 |
| 0.03810714 | 5.712417208 | |
| 0.019984511 | 33.32345909 | |
| 0.020451241 | 36.31601725 | 0.010081468 |
| 0.030795641 | 11.67418496 | |
| 0.017973397 | 35.62636494 | |
| 0.018882637 | 37.82489969 | 0.036507775 |
| 0.002888025 | 111.5882306 | |
| 0.002374001 | 128.6031261 | |
| 0.001034184 | 255.5243197 | 0.03552477 |
| 0.001165094 | 200.2197564 | 0.035132167 |
| 0.008307219 | 37.00721075 | |
| 0.00513847 | 51.28714501 | |
| 0.00282037 | 95.32830788 | 0.122636056 |

| | | |
|---|---|---|
| 0.00266963 | 82.27007015 | 0.093395738 |
| 0.016635631 | 22.99060476 | |
| 0.022584786 | 89.40823182 | |
| 0.013286746 | 76.61516523 | |
| 0.024028198 | 88.43742286 | 0.222198804 |
| 0.007577443 | 6.185031554 | 0.103190615 |
| 0.00800248 | 6.198758366 | 0.084839589 |
| 0.020190951 | 3.520222858 | 0.367341522 |
| 0.028611074 | 3.022320897 | 0.397066233 |

The yellow values in this column were used in Figure S2F

| WT raw flotation ratio | 2K-E concentration back-calculated (μM) | 2K-E concentration (μM) |
|---|---|---|
| 3.028032834 | 0.126700729 | 0.065077827 |
| 4.012241602 | 0.126700729 | 0.051815981 |
| 3.489886166 | 0.126700729 | 0.059633047 |
| 5.102717404 | 0.126700729 | 0.04846415 |
| 2.426620289 | 0.055748321 | 0.038593372 |
| 4.741470417 | 0.055748321 | 0.038085183 |
| 3.467373868 | 0.055748321 | 0.040094136 |
| 5.480623155 | 0.055748321 | 0.039870035 |
| 2.05573808 | 0.055748321 | 0.064045674 |
| 3.398424693 | 0.055748321 | 0.064254598 |
| 2.084019145 | 0.055748321 | 0.08230153 |
| 3.665005603 | 0.055748321 | 0.078155433 |
| 4.429209968 | 0.063350365 | 0.234573339 |
| 7.596447218 | 0.063350365 | 0.237104063 |
| 5.737604007 | 0.063350365 | 0.24524518 |
| 8.61586839 | 0.063350365 | 0.220045469 |
| 4.408919282 | 0.063350365 | 0.052585983 |
| 6.288787509 | 0.063350365 | 0.02326038 |
| 7.107279503 | 0.063350365 | 0.014568921 |
| 5.907208804 | 0.063350365 | 0.035098969 |
| 5.766033759 | 0.063350365 | 0.021811078 |
| 6.101763032 | 0.063350365 | 0.015976282 |
| 2.43634079 | 0.063350365 | 0.347561266 |
| 4.567341067 | 0.063350365 | 0.44420769 |
| 3.144118946 | 0.063350365 | 0.463795154 |
| 5.290747238 | 0.063350365 | 0.55399829 |
| 2.933338261 | 0.063350365 | 0.321012226 |
| 11.23683236 | 0.063350365 | 1.106750726 |
| 4.080654428 | 0.063350365 | 0.324290009 |
| 11.52584221 | 0.063350365 | 0.438435592 |
| 2.284966883 | 0.063350365 | 0.036654296 |
| 13.32938364 | 0.063350365 | 0.019314992 |
| 14.5264069 | 0.063350365 | 0.018856695 |
| 4.669673983 | 0.063350365 | 0.029722524 |
| 14.25054597 | 0.063350365 | 0.017669532 |
| 15.12995988 | 0.063350365 | 0.017487977 |
| 44.63529223 | 0.076020438 | 0.012563989 |
| 51.44125044 | 0.076020438 | 0.008566082 |
| 102.2097279 | 0.076020438 | 0.003938231 |
| 80.08790257 | 0.076020438 | 0.002906461 |
| 14.8028843 | 0.076020438 | 0.011543522 |
| 20.514858 | 0.076020438 | 0.007608163 |
| 38.13132315 | 0.076020438 | 0.00538002 |

| | | |
|---|---|---|
| 32.90802806 | 0.076020438 | 0.004897096 |
| 9.196241903 | 0.050680292 | -0.003457716 |
| 35.76329273 | 0.050680292 | 0.014612778 |
| 30.64606609 | 0.050680292 | 0.009457874 |
| 35.37496914 | 0.050680292 | 0.012100105 |
| 2.474012622 | 0.076020438 | 0.02064038 |
| 2.479503346 | 0.076020438 | 0.022794201 |
| 1.408089143 | 0.050680292 | 0.016475196 |
| 1.208928359 | 0.050680292 | 0.01811668 |

| 2K-E binding percentage (same blot) | 2K-E raw flotation ratio | WT concentration (computed) |
|---|---|---|
| 4.330683438 | 1.732273375 | 0.512438393 |
| 5.026580428 | 2.010632171 | 0.407370896 |
| 5.460156619 | 2.184062648 | 0.54147742 |
| 6.218021379 | 2.487208551 | 0.397601401 |
| 5.782731184 | 2.313092474 | 0.857201035 |
| 6.259575474 | 2.50383019 | 0.892351445 |
| 6.457054312 | 2.582821725 | 0.991717014 |
| 7.331432718 | 2.932573087 | 0.956834545 |
| 5.764640043 | 2.305856017 | 1.56582451 |
| 5.592940759 | 2.237176303 | 1.623458961 |
| 5.4659233 | 2.18636932 | 2.19556663 |
| 6.838162907 | 2.735265163 | 1.964252797 |
| 4.125882429 | 1.650352972 | 3.32267498 |
| 4.062381808 | 1.624952723 | 3.533857223 |
| 4.728180916 | 1.891272366 | 3.496814171 |
| 4.578002803 | 1.831201121 | 3.026300788 |
| 4.17595948 | 1.670383792 | 2.56692321 |
| 3.665942897 | 1.466377159 | 1.070826759 |
| 4.110590767 | 1.644236307 | 0.696608855 |
| 5.463605263 | 2.185442105 | 1.722512573 |
| 4.999833544 | 1.999933418 | 1.186193064 |
| 5.375682648 | 2.150273059 | 0.797622268 |
| 8.138087502 | 3.255235001 | 13.27370255 |
| 9.289831178 | 3.715932471 | 16.75853514 |
| 8.965717373 | 3.586286949 | 17.91707197 |
| 11.33059619 | 4.532238476 | 22.37386403 |
| -0.593009287 | -0.237203715 | 17.57126832 |
| -0.814821949 | -0.325928779 | 59.63413085 |
| 1.475844234 | 0.590337694 | 17.81329871 |
| 0.524886715 | 0.209954686 | 27.14970542 |
| 0.738047584 | 0.295219034 | 0.889166609 |
| 0.339838343 | 0.135935337 | 0.466305266 |
| 0.383775348 | 0.153510139 | 0.477195625 |
| 1.49282472 | 0.597129888 | 0.718564955 |
| 1.333926998 | 0.533570799 | 0.419379272 |
| 1.465821566 | 0.586328626 | 0.440594868 |
| 7.026752624 | 2.81070105 | 0.067387246 |
| 9.254730116 | 3.701892046 | 0.055393358 |
| 8.601247515 | 3.440499006 | 0.02413096 |
| 10.73770776 | 4.295083106 | 0.027185515 |
| 8.92802387 | 3.571209548 | 0.19383511 |
| 11.148698 | 4.4594792 | 0.119897627 |
| 9.074199251 | 3.6296797 | 0.065808645 |

| | | |
|---|---|---|
| 7.990227321 | 3.196090928 | 0.06229136 |
| -75.11047359 | -30.04418944 | 0.38816472 |
| 19.51333438 | 7.805333753 | 0.526978345 |
| 33.76142384 | 13.50456954 | 0.310024085 |
| 27.77696078 | 11.11078431 | 0.560657961 |
| 0.638237211 | 0.255294884 | 0.176807007 |
| 0.52590011 | 0.210360044 | 0.186724539 |
| 1.293124023 | 0.517249609 | 0.314081454 |
| 1.200061573 | 0.480024629 | 0.445061153 |
| | | 1.080432145 |

| 2K-E concentration (computed) |
|---|
| 0.911089572 |
| 0.725423736 |
| 0.834862654 |
| 0.678498107 |
| 1.227970915 |
| 1.211801274 |
| 1.275722502 |
| 1.26859202 |
| 2.037816886 |
| 2.044464483 |
| 2.618685042 |
| 2.48676377 |
| 6.568053488 |
| 6.638913761 |
| 6.866865052 |
| 6.161273127 |
| 1.472407528 |
| 0.651290645 |
| 0.407929782 |
| 0.982771136 |
| 0.610710174 |
| 0.447335904 |
| 9.731715441 |
| 12.43781533 |
| 12.98626431 |
| 15.51195212 |
| 8.988342339 |
| 30.98902033 |
| 9.08012024 |
| 12.27619658 |
| 1.026320277 |
| 0.54081979 |
| 0.527987455 |
| 0.832230686 |
| 0.494746897 |
| 0.489663346 |
| 0.293159739 |
| 0.19987525 |
| 0.091892062 |
| 0.06781742 |
| 0.269348848 |
| 0.177523804 |
| 0.125533809 |

| WT concentration (µM) | WT raw flotation ratio |
|---|---|
| 0.036602742 | 3.028032834 |
| 0.029097921 | 4.012241602 |
| 0.038676959 | 3.489886166 |
| 0.0284001 | 5.102717404 |
| 0.061228645 | 2.426620289 |
| 0.063739389 | 4.741470417 |
| 0.07083693 | 3.467373868 |
| 0.068345325 | 5.480623155 |
| 0.111844608 | 2.05573808 |
| 0.115961354 | 3.398424693 |
| 0.156826188 | 2.084019145 |
| 0.140303771 | 3.665005603 |
| 0.142400356 | 4.429209968 |
| 0.151451024 | 7.596447218 |
| 0.149863464 | 5.737604007 |
| 0.129698605 | 8.61586839 |
| 0.110010995 | 4.408919282 |
| 0.045892575 | 6.288787509 |
| 0.029854665 | 7.107279503 |
| 0.073821967 | 5.907208804 |
| 0.050836846 | 5.766033759 |
| 0.034183811 | 6.101763032 |

| | |
|---|---|
| 0.007577443 | 2.474012622 |
| 0.00800248 | 2.479503346 |
| 0.020190951 | 1.408089143 |
| 0.028611074 | 1.208928359 |

The orange values above were used in Figure S2F

| |
|---|
| 0.114265563 |
| -0.12102007 |
| 0.511447246 |
| 0.331025575 |
| 0.423503668 |
| 0.481608857 |
| 0.531864701 |
| 0.576631857 |
| 0.634083808 |
| 1.773810212 |

| 2K-E concentration (µM) | 2K-E raw flotation ratio | WT raw fraction bound | 2K-E fraction bound |
|---|---|---|---|
| 0.065077827 | 1.732273375 | 0.433008695 | 0.247715093 |
| 0.051815981 | 2.010632171 | 0.573750549 | 0.2875204 |
| 0.059633047 | 2.184062648 | 0.499053722 | 0.312320959 |
| 0.04846415 | 2.487208551 | 0.729688589 | 0.355670823 |
| 0.038593372 | 2.313092474 | 0.347006701 | 0.330772224 |
| 0.038085183 | 2.50383019 | 0.67803027 | 0.358047717 |
| 0.040094136 | 2.582821725 | 0.495834463 | 0.369343507 |
| 0.039870035 | 2.932573087 | 0.783729111 | 0.419357951 |
| 0.064045674 | 2.305856017 | 0.293970545 | 0.32973741 |
| 0.064254598 | 2.237176303 | 0.485974731 | 0.319916211 |
| 0.08230153 | 2.18636932 | 0.298014738 | 0.312650813 |
| 0.078155433 | 2.735265163 | 0.524095801 | 0.391142918 |
| 0.234573339 | 1.650352972 | 0.633377025 | 0.236000475 |
| 0.237104063 | 1.624952723 | 1.086291952 | 0.232368239 |
| 0.24524518 | 1.891272366 | 0.820477373 | 0.270451948 |
| 0.220045469 | 1.831201121 | 1.23206918 | 0.26186176 |
| 0.052585983 | 1.670383792 | 0.630475457 | 0.238864882 |
| 0.02326038 | 1.466377159 | 0.899296614 | 0.209691934 |
| 0.014568921 | 1.644236307 | 1.016340969 | 0.235125792 |
| 0.035098969 | 2.185442105 | 0.844730859 | 0.312518221 |
| 0.021811078 | 1.999933418 | 0.824542828 | 0.285990479 |
| 0.015976282 | 2.150273059 | 0.872552114 | 0.307489047 |
| 0.02064038 | 0.255294884 | 0.353783805 | 0.036507168 |
| 0.022794201 | 0.210360044 | 0.354568979 | 0.030081486 |
| 0.016475196 | 0.517249609 | 0.201356747 | 0.073966694 |
| 0.01811668 | 0.480024629 | 0.172876755 | 0.068643522 |
